# A sensory approach to turbidity: How sources and levels shape aquatic light environments and fish visual ecology

**DOI:** 10.1101/2024.10.10.617539

**Authors:** Anna Garcia, Adelaide Sibeaux, Theresa Burt de Perera, Cait Newport

## Abstract

Turbidity is a ubiquitous source of sensory pollution that is likely to impact the appearance of the visual stimuli that animals rely on for survival and reproduction. Understanding how different turbidity sources impact the appearance of the ambient light environment is the foundational first step towards predicting whether and how animals will cope with the global increases in the severity and frequency of high turbidity events caused by anthropogenic disturbance. Here, we measured how four common turbidity sources (algae, bentonite, calcium carbonate, and kaolin), and variable turbidity levels, changed the appearance of the ambient light environment. We measured total number of photons (luminance), hue, chroma, and image contrast, and we evaluated the effect of each turbidity source and level on settling rate, pH, and KH. Both turbidity source and turbidity level impacted the appearance of the ambient light environment. With increasing turbidity level, calcium carbonate and kaolin increased luminance while algae decreased luminance, bentonite caused the greatest change in hue, and algae caused the greatest change in chroma. This demonstrates that the impacts of different turbidity sources on the ambient light environment are not uniform, giving a potential explanation for the discrepancies between studies on the effects of turbidity on fish behaviour. Consideration of the effect of specific turbidity sources on ambient light is crucial for the design of experiments that seek to investigate how changes in turbidity impact the perception of important visual information, which underpins the survival and reproductive success of aquatic organisms around the world.

**Summary Statement:** Different turbidity sources and levels uniquely affect aquatic light environments and fish visual perception, emphasizing the need for these factors when interpreting fish behaviour and the ecological consequences of turbidity.

## Introduction

Turbidity refers to organic or inorganic particles in water which can alter both water clarity and quality. While the impacts of turbidity on aquatic ecosystems are likely to be wide ranging, its most visible impact is on the light environment. Particles in the water scatter and absorb light and can significantly alter the appearance of visual features, potentially making visual signals more difficult to detect or identify. Turbidity levels therefore play an important role in shaping the visual environment of aquatic species. Turbidity occurs naturally to some degree in all aquatic ecosystems and fluctuates over both short and long-time scales, suggesting that visually reliant animals likely have some mechanisms to cope with the associated changing visual conditions. However, natural turbidity conditions are being increasingly altered through human activities (Burke, Reytar et al. 2011, Seers and Shears 2015, Cartwright, Fearns et al. 2021), changing both the concentration of turbidity as well as the source. For example, more frequent flooding in Australia is causing plumes of turbidity that are visible from space to drain onto the Great Barrier Reef (Malthus, Lehmann et al. 2019), and sewage and chemical discharges in British rivers and coasts are turning normally clear waters to an opaque brown (Bojko, Lipp et al. 2020, Love, Crooks et al. 2020, Albini, Lester et al. 2023). For example, more frequent flooding in Australia is causing plumes of turbidity that are visible from space to drain onto the Great Barrier Reef (Malthus, Lehmann et al. 2019) and sewage and chemical discharges in British rivers and coasts are turning normally clear waters to an opaque brown (Bojko, Lipp et al. 2020, Love, Crooks et al. 2020, Albini, Lester et al. 2023). It is not just turbidity levels that are changing; in some ecosystems, the sources of turbidity have also shifted, leading to the presences of multiple sources in some ecosystems. For example, in addition to the inorganic turbidity associated with coastal run-off, the Great Barrier Reef is likely to experience an increase in organic turbidity as bleaching events can lead to eventual algae blooms (Fukunaga, Burns et al. 2022). In the UK, nutrient runoff from intensive agricultural practices can accelerate the export of carbon and other organic nutrients resulting in algal blooms (Glendell and Brazier 2014). There is some evidence that turbidity can negatively impact fish. For example, studies have shown that turbidity can alter species richness of coral reef fish (Moustaka, Langlois et al. 2018) and disrupt fish community structures in shallow lakes (Hayes, Rutledge et al. 1992). Additionally, turbidity has also been linked to increased gill epithelium thickness in clownfish larvae (Amphiprion percula) (Hess, Wenger et al. 2015), reduced foraging success and growth rate in smallmouth bass (Micropterus dolomieu) (Sweka and Hartman 2011), decreased survival of shed larvae (Alosa sapidissima) (Auld and Schubel 1978), and altered metabolism in rainbow trout (Berli, Gilbert et al. 2014). Furthermore, turbidity can contribute to eutrophication and anoxia (Smith 2003, Budria and Candolin 2015, Cumming and Herbert 2016), impairment of metabolic reduced physiological condition (Budria and Candolin 2015, Hasenbein, Fangue et al. 2016), and negatively affect spawning success (Burkhead and Jelks 2001, Sutherland and Meyer 2007). While it seems likely that the effects of turbidity could be wide-ranging, its impact on visually-guided fish behaviour has received less attention, leaving it unclear whether turbidity presents a serious threat to fish populations.

Turbidity has been shown to significantly alter some visually-guided behaviours in fish, including foraging (Lunt and Smee 2015) and searching (Newport, Padget et al. 2021), collective behaviour (Chamberlain and Ioannou 2019), predator-prey interactions (Ortega, Figueiredo et al. 2020), and swimming rates (Hildebrandt and Parsons 2016). For guppies (*Poecilia reticulata*), increased turbidity causes individuals to be less active, more solitary (Borner, Krause et al. 2015), and more vulnerable to predation (Kimbell and Morrell 2015). But some species acclimate to changing visual conditions; the same guppy species raised in turbid water, display a shift in spectral sensitivity and an increase in activity (Ehlman, Sandkam et al. 2015). African cichlids (*Pseudocrenilabrus multicolour*) reared in turbid conditions exhibited an ontogenetic shift in the relative size of their eyes and brains, with young fish raised in turbid water developing relatively larger eyes, and older fish developing relatively larger brains compared to those raised in clear water (Tiarks, Gray et al. 2024). Studies exploring the impacts of turbidity on the expression of sexually selected traits have revealed complex effects on both sexes, making it difficult to predict the outcomes of such interactions (Järvenpää and Lindström 2004, Järvenpää and Lindström 2011, Sundin, Rosenqvist et al. 2016, Sundin, Aronsen et al. 2017, Järvenpää, Pauli et al. 2019). For example, turbidity interferes with the visual inspection of conspicuous red nuptial coloration by female three-spined sticklebacks (*Gasterosteus aculeatus*), altering both female mate choice preference, and male courtship behaviour (Wong, Candolin et al. 2007, Tuomainen and Candolin 2011, Candolin and Jensen 2021). However, there is also evidence that turbidity can have no impact on sexual selection behaviour, as shown studies on yellow perch (*Perca flavescens*), wreakfish (*Cynoscion regalis*), white perch (*Morone americana*), and tessellated darter (*Etheostoma olmstedi*), (Grecay and Targett 1996, Shoji, North et al. 2005, Pangle, Malinich et al. 2012, Kellogg and Leipzig-Scott 2017). In sum, the effects of turbidity on fish are neither unidirectional nor easily predicted, and the evidence suggests that not all fish species or behaviours are equally affected by turbidity (Rodrigues, Ortega et al. 2023).

The disparity in the types and magnitude of effects observed between studies may also arise from methodological differences, particularly the lack of consideration for how specific turbidity conditions alter the light environment and, consequently the transmission of visual information that experimenters aim to manipulate. Differences in turbidity particle size, shape, composition, and concentration influence the scattering of light. Small particles whose radii are less than 10% of the wavelength of light obey Rayleigh scattering. In this case, the scattering of light is strongly wavelength and intensity-dependent, and shorter wavelengths (e.g. violet) are scattered more than longer wavelengths (e.g. red) (Johnsen 2012). Larger particles that are comparable to or larger, than the wavelength of light, follow Mie scattering which is relatively wavelength-independent (Bohren and Huffman 2008). Both forms of scattering are dependent on the total light intensity (i.e. number of photons available) whereby smaller particles scatter more light than larger particles. Moreover, light absorption is highly dependent on the properties of the turbidity source. For example, photopigments like chlorophylls in algal blooms selectively absorb light in the 440-475 nm and 630-675 nm ranges (Bricaud, Claustre et al. 2004). On the other hand, mineral and sediment-derived turbidity sources absorb in the shorter 400-550 nm wavelength range (Babin, Morel et al. 2003). Despite the source-dependent effects of turbidity on the transmission of important visual information, previous research seldom considers how different types of turbidity may influence experimental outcomes (but see (Radke and Gaupisch 2005, Li, Zhang et al. 2013, Cano-Rocabayera, Vargas-Amengual et al. 2020, Figueiredo, Granzotti et al. 2020)).

To understand how different sources and levels of turbidity have been addressed in the literature, we conducted a review of existing studies. We identified 200 papers that explore the impact of turbidity on a range of fish behaviours including anti-predator, foraging efficiency, movement, sexual selection, social/collective behaviour (Fig. 1). In this review, we also included papers assessing the impact of different turbidity sources on fish physiological conditions and biodiversity as these parameters are directly related to fish behavioural ecology. See supplementary materials S1 for details of the literature search process. We found that the turbidity sources used in experiments can be broadly categorised into eight groups (Fig. 1) and that natural sediment, bentonite clay, and algae/phytoplankton were the most used turbidity sources. We also found that the use of a particular turbidity source is not field- or species-specific. For example, the studies examining the impacts of turbidity on foraging efficiency, used all eight of our turbidity groups. Similarly, different studies involving the same species, Trinidadian guppies (*Poecilia reticulata),* used different turbidity source and concentrations. While Borner, Krause et al. (2015) explored the effects of turbidity on social dynamics using Kaolin, a white clay with a fine particle size (>1 µm), at concentrations of up to 1000 Nephelometric Turbidity Units (NTU), Kimbell and Morrell (2015) tested behavioural responses to predation using Bentonite, a typically reddish clay with even smaller particles (>1 µm), at levels of up to 200 NTU. Interestingly, the two groups cite very different reasons for their choice of turbidity source. While Borner, Krause et al. (2015) made their selection based on practical considerations (kaolin was thought to stay suspended for longer), Kimbell and Morrell (2015) chose a source that was ecologically relevant. While both reasons are valid, surprisingly neither of author considered how each turbidity source would alter the appearance or detectability of conspecifics or predators, which was at the heart of the behaviours being tested in their studies. This is a common weakness of many studies on this topic including our own (Newport, Padget et al. 2021).

**Figure 1.**
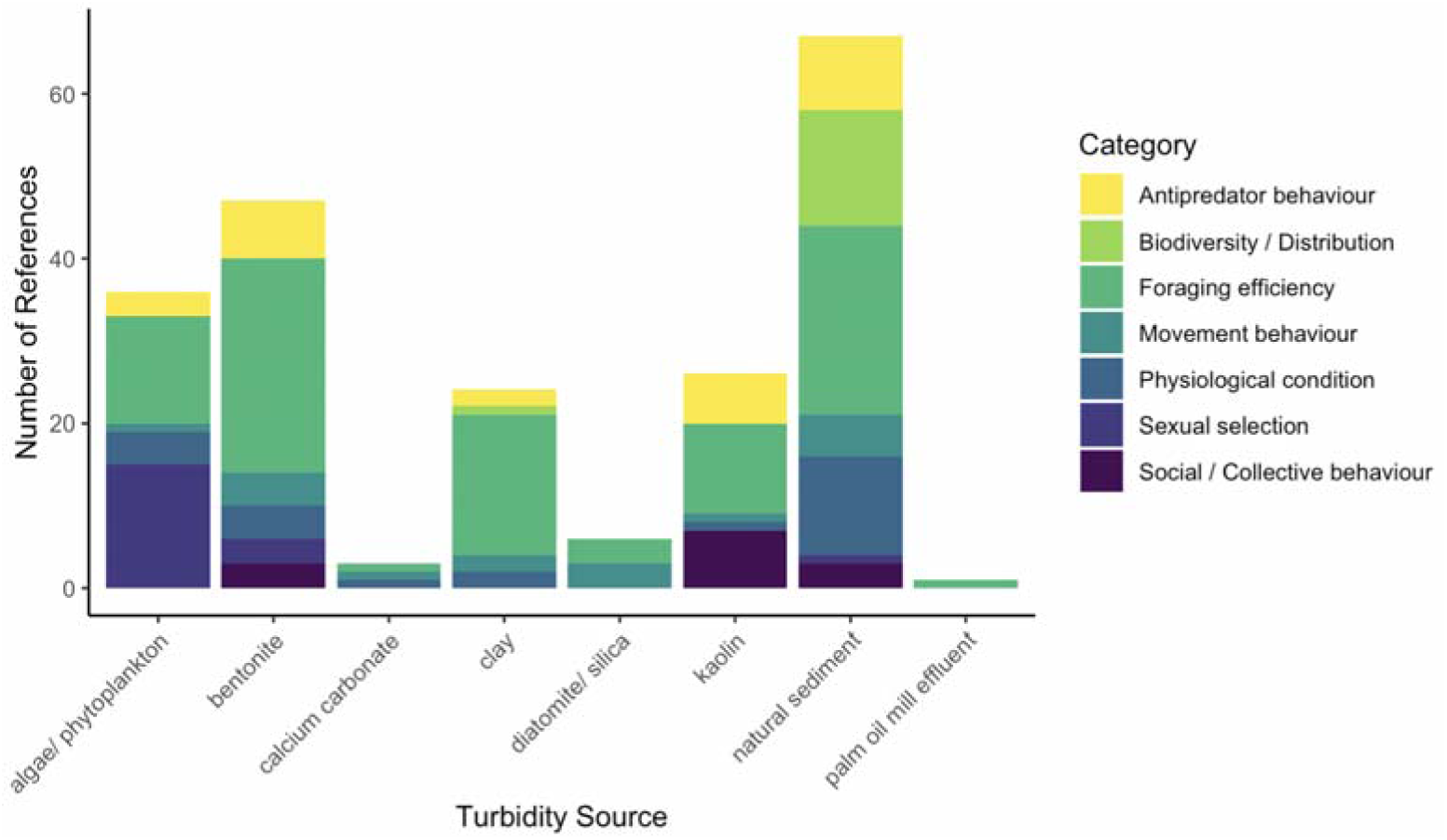
The frequency of use of different turbidity sources in 200 scientific papers exploring the impact of turbidity on fish behaviour, physiological condition, and biodiversity. Papers are grouped by turbidit source. “Clay” and “natural sediment” are often sourced directly from the study site, with their specific composition unknown. More specific turbidity sources, such as bentonite, were grouped separately. Within each turbidity source, papers are categorized by research focus. Studies using multiple turbidit sources or response categories were counted separately for each.

The aim of this paper is to describe how different sources and levels of turbidity impact the ambient light environment and conspicuousness of visual features, as well as some practical considerations associated with using turbidity in laboratory behavioural experiments. We hope this will inspire other researchers to consider turbidity sources carefully in their experimental design and empower them to select the appropriate turbidity sources based on their experimental investigation and design.

The ability of an animal to perceive visual features depends not only on its visual system, but also on four additional optical factors that determine the light available to its eyes: 1) the light environment, 2) the amount of light reflected by the stimulus, 3) the light transmitted between the feature and the eye, 4) and the degree of veiling light (light scattered from particles directly into the observer’s eye) (Endler 1990). These four factors determine the amount of light that reaches the eye and marks the beginning of the visual processing pathway. This study focuses on how turbidity affects the first factor – the light environment – aiming to provide an in-depth understanding of how turbidity influences the appearance of visual features within the environment. Absolute irradiance is commonly used to describe the light environment. It is a measure of the number of photons per wavelength within a light environment and is visualized as a spectral curve. Changes to the shape of this curve indicate changes in the light environment. For example, shifts in the position of the curve’s peaks indicate changes in colour composition, while flattening peaks shows reduced light availability and lower spectral saturation. The shape of the curve can be described using three characteristics: luminance, hue, and chroma (Endler 1990). Luminance is calculated from the area under the spectral curve, hue is determined by the wavelength range at which the number of photons peaks, and chroma represents the difference between the lowest and highest peaks across all wavelengths (see Endler 1990 for mathematical formulas). These characteristics describe how the available light will influence the appearance of visual features within the light environment and are independent any species’ specific visual system. The specific perception and interpretation of visual signals will depend on the visual system of individual species, but the raw optical input dictates what is possible and serves as a basis for visual modelling. Therefore, this study provides a critical foundation for predicting how turbidity might alter the appearance of visual stimuli by examining the effects of different turbidity levels and sources on the light environment.

## Methods

We tested how four sources of turbidity commonly used in behavioural experiments – algae, bentonite, kaolin, calcium carbonate – alters the light environment. We specifically measured changes in absolute irradiance, and image contrast. Changes to luminance, hue, and chroma were calculated from absolute irradiance measurements. We also measured turbidity settling rate, as well as changes in pH and water hardness (calcium carbonate concentration, KH) over time to assess the usability of each turbidity source in behavioural experiments with fish. The latter two were measurements help determine whether dissolving particles significantly alter water chemistry, potentially exceeding the healthy limits for fish. We tested turbidity levels between 0 and 100 Nephelometric Turbidity Units (NTU) which reflect those commonly found in nature (Cyrus and Blaber 1992, Miranda 2011, Li, Zhang et al. 2013, Macdonald, Ridd et al. 2013, Ehlman, Sandkam et al. 2015) and used in experiments (Ajemian, Sohel et al. 2015, Borner, Krause et al. 2015, Kimbell and Morrell 2015, Newport, Padget et al. 2021).

### General Apparatus

Experiments were conducted in a temperature-controlled laboratory (21.2 ± 0.587 °C; mean ± standard deviation) with artificial fluorescent lighting that followed a 12-hour light/dark cycle (Clean A-MP LED6000-840 M600Q LDO, total power 46.3 W LED, Zumtobel). A glass tank with dimensions 84 cm x 30 cm x 57.5 cm (l x h x w) (Fig. 2) was filled to a height of 28 cm (volume = 135 litres). RO water was used to provide a standardised, replicable mineral composition of the water. Opaque white PVC sheets were secured outside the tank against the walls and under the base to standardize the source of incoming light. A tripod was secured inside the tank to the bottom to hold either 1) an optic fibre cable at 15 cm from the bottom of the tank allowing to measure total photons and spectral composition of the light in the water, or 2) a GoPro (HERO8 Black, wide lens, photo, standard output, zoom x1. GoPro, Inc.) for image contrast measurements. The GoPro was attached to the tripod facing the image standard, with its lens at a height of 15 cm.

**Figure 2.**
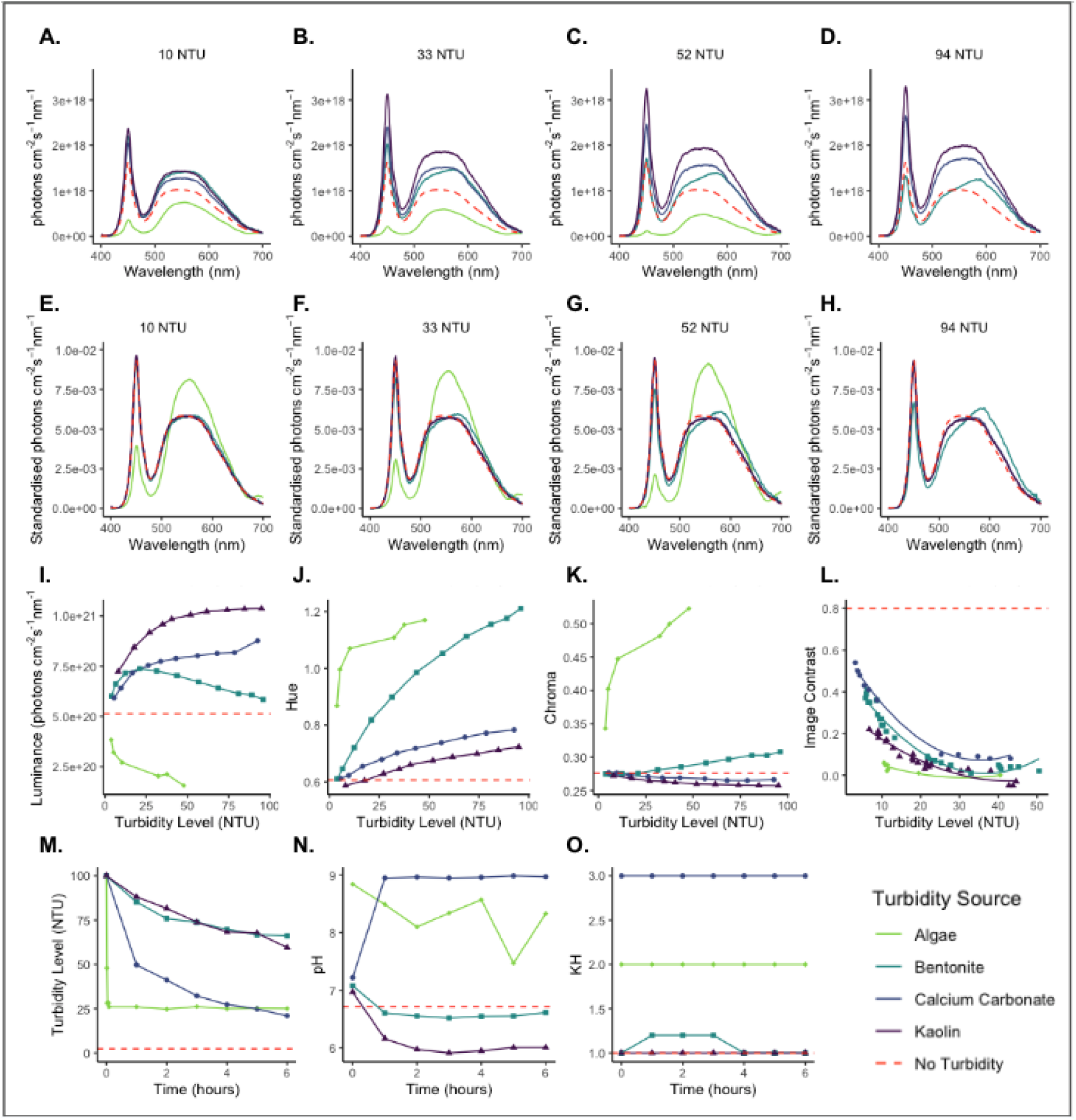
**(A-H) Effect of turbidity on absolute irradiance:** A-D show raw light spectra and E-H show standardised spectra with the area under the curve set to one. (**I-L) Effect of turbidity on luminance, hue, chroma, and image contrast.** Individual points indicate observed values. In panel L, lines represent the best-fitting model, which shows a significantly non-linear decrease in image contrast with increasing turbidity. In panels I-K, algae were only measured up to 50 NTU as we could not achieve a turbidity level above this under our growing conditions. In panel L, contrast measurements were taken from 0-120 NTU, but only 0-50 NTU are displayed as contrast values > 50 NTU approached zero. (**M-O) Turbidity settling rate, and the effects of turbidity on pH and water hardness (KH) over time.** For visualisation purposes, settling rate, pH, and KH data points are presented as the average percentage change in each value between time points, then re-transformed to fit the original scale.

Directly across from the tripod, a laminated (Office Depot laminating pouches, matt, 75 µm) image standard printed on paper (Canon, Matte Photo Paper), was attached to the back wall with Blu Tack (Bostick Smart Adhesives) (see Supplementary Fig. 1, S2 for an illustration of the apparatus). We included an image standard to ensure that colour was present in the tank, allowing us to measure changes in colour caused by turbidity. The image standard was designed with colours selected based on the cone sensitivity maxima of a coral reef fish species *(Rhinecanthus* aculeatus) and goldfish, ensuring the colours are visible to two species that inhabit very different visual environments. The orange (Hex colour: #ff6b00) and blue (#0035ff) squares match the goldfish cone sensitivity peaks at 623 nm and 447 nm, respectively (Palacios, Varela et al. 1998). The green (#56ff00) and purple (#7900e6) squares correspond to the *R. aculeatus* cone sensitivity peaks at 528 nm and 413 nm, respectively (Cheney, Newport et al. 2013). The width of the black and white stripes was chosen to be discriminable from a viewing distance of 1 m for both species (Neumeyer 2003, Cheney, Hudson et al. 2022).

### Turbidity preparation

Turbidity was measured in Nephelometric Turbidity Units (NTU) using a turbidimeter (2100Q Portable Turbidimeter, Hach). For all turbidity measurements, three 15 ml water samples were collected and averaged. Prior to measurement in the turbidimeter, samples were inverted several times to ensure particles remained suspended. Inorganic turbidity sources were purchased in dry powder form (bentonite clay: Fullers Earth, Trustleaf, Cambridgeshire UK; calcium carbonate: Intra Laboratories, Plymouth, UK; kaolin clay: Trustleaf, Cambridgeshire UK). Each turbidity source was mixed by hand with small additions of RO water and sieved through a hand net. This step was required to prevent particles from clumping and causing uneven particle distribution. A stock solution of known concentration was created and used to gradually increase turbidity within the experimental tank to the desired concentration of 100 NTU. The amount of inorganic turbidity (bentonite: 5.04 × 10^−2^ grams per litre (g/L); calcium carbonate: 2.22 × 10-2 g/L; kaolin: 1.19 × 10^−2^ g/L).

Live freshwater algae, *Scenedesmus quadricauda* (Blades Biological Ltd) was purchased and grown over a 12-week period to produce 135L of water at 100 NTU. *S. quadricauda* was chosen because of its fast growth rate and preferred temperature range which matched that of our lab. To start, 240 mL of culture starter was added to the experimental tank which was half filled (approximately 70L), along with 80 mL of Alga-Gro growth medium (Blades Biological Ltd). A glass lid was placed loosely over the tank to reduce contamination, and a grow light (9 hour on/ 15 hour off) was attached over the tank (40 cm above tank). The mixture was gently stirred by hand every other day to increase aeration and reduce algal clumping. Over the four weeks of growth, the maximum turbidity level reached was 9 NTU so the tank was moved outside to a greenhouse on February 6th, 2024. A commercial aquatic plant fertiliser (AquaDesign PLANT+ 1000ML Aquarium Plant “all in one” Complete Liquid Fertiliser) was added and stirred weekly, and the algae was left to grow for a further 8 weeks until it had reached an average turbidity level measurement of 50 NTU. Algal turbidity levels plateaued at 12 weeks, so algae turbidity measurements were reduced from 0-100 to 0-52 NTU for this turbidity source.

### Light measurements

The side-welling absolute irradiance was measured with an Ocean Insights Flame-S-UV-VIS spectrometer with a 400 µm fibre and a cosine corrector (CC-3 Spectralon optical diffuser, Ocean Insight). The fibre optic cable was held in place with a 3D printed mount and measurements were taken from the centre of the tank, 25 cm from the image standard, 15 cm above the bottom of the tank, with the cosine corrector pointing towards the image standard.

### Experimental procedure

The following procedure was repeated for each turbidity source. The experimental tank was first filled with RO water and the tripod secured to the bottom of the tank 50 cm from the image standard (Supplementary Fig. S2). A wavemaker (TMC Reef Flow 2.0 16000 DC Wavemaker; intensity setting: 2, mode: 1) was used throughout tests with inorganic turbidity sources to ensure particles remained suspended. The wavemaker was removed for tests with algae to prevent damage to the algal cells. Prior to adding any turbidity, baseline measurements and absolute irradiance were taken in clear water. Approximately 100 mL of turbidity solution was then added to the experimental tank to achieve a turbidity level between 0-5 NTU. The two measurements (NTU and absolute irradiance) were repeated. This process continued, increasing the turbidity in increments of 10 NTU, up to a maximum of 100 NTU. Once the final measurement had been taken, the tank was emptied, cleaned, and refilled with RO water to test the next turbidity source.

A similar procedure was used to measure image contrast, but the camera was mounted on the tripod instead of the fibre optic cable. The camera was positioned 15 cm from the image standard, at an optimal distance determined by pilot experiments to best capture image contrast at different turbidity levels. As previously described, baseline measurements were taken (i.e. turbidity level, contrast photo) in clear water and repeated as turbidity was gradually increased. Once a turbidity of 100 NTU was reached, measurements were taken and then repeated every hour over a total of six hours. The final turbidity level measurement of a trial day determined the starting turbidity level of the next trial day. The tank was emptied, cleaned, and refilled and this protocol repeated the next trial day at the next starting turbidity level. This process was repeated until the turbidity level dropped to 10 NTU by the final hour of an experimental day. The number of trial days required to reach this criterion varied for each turbidity source (algae: n = 1 trial day, bentonite: n = 5 trial days or 35 measurements; calcium carbonate: n = 2 trial days or 14 measurements; kaolin clay: n = 5 trial days or 35 measurements).

Finally, the procedure was repeated to measure how turbidity level, pH and water hardness changed over a six-hour period. The experimental tank was initially set to a turbidity of 100 NTU measured. The following four measurements were taken every hour: 1) turbidity, 2) pH (Chauvin Arnoux waterproof C.A 10001 pH metre), 3), temperature, and 4) KH (API KH test, API Fish Care).

### Analysis

#### Data Treatment

##### Absolute irradiance

The number of photons at each wavelength (photons cm^−2^ s^−1^ nm^−1)^ were extracted from raw spectrometer measurements using OceanView software (Ocean Insight; Version 2.0.8). Absolute irradiance was plotted as a spectral curve for each experimental condition and allowed the shape of the curves to be compared. Spectral curves were also standardized so that the area under each curve equals 1, effectively normalizing the overall light levels. Four turbidity level categories were chosen to compare spectral curve shape characteristics: 1) 10 NTU (±3 NTU), 2) 33 NTU (±2 NTU), 3) 52 NTU (±4.5 NTU), and 4) 94 NTU (±2 NTU). The final category did not include spectral measurements for algae.

Absolute irradiance spectrum values were used to calculate luminance, hue and chroma (see Endler 1990). Luminance refers to the total number of photons and is the achromatic property of the light environment. Hue and chroma are determined by the relative number of photons in different wavebands of the light spectrum and are the chromatic properties of the light environment. Hue is determined by the wavelength region with the steepest slope in the spectra, as well as whether the slope is positive or negative. The hue of the ambient light environment is the category of colour, calculated using the segment classification method (Endler, 1990). This method is used to assess changes to the chromatic properties of hue and chroma independently of a specific visual system while still preserving the fundamental processes of ‘opponency’ and ‘lateral inhibition’ common to many invertebrate and invertebrate species. This produces a two-dimensional colour space with the opponent pairs of long to medium-short wavelengths on the vertical axis, and short to medium-long wavelengths on the horizontal axis. Hue is calculated as the clockwise angle from the vertical axis about the origin, where the origin represents full spectrum white light. This value indicates the waveband in the spectra with the greatest number of photons. Chroma is the chromatic saturation of the light environment and is determined by the magnitude of the difference in the number of photons between different parts of the spectrum and the maximum slope of the spectra. A spectrum with more photons in one waveband compared to its opponent pair waveband, will have a higher chroma than a spectrum with equal/similar numbers of photons within opponent pairs.

#### Image Contrast

The image contrast of the photos of the image standard was calculated using Michelson contrast (Michelson 1927), which measures the contrast gradient of a sinusoidal pattern on a scale of 0 to 1. Michelson contrast gives the relation between the spread and the sum of two luminance values and is commonly used in signal processing theory to determine the quality of a signal relative to its noise level (e.g. scattered light) (Cronin, Johnsen et al. 2017, Hunter, Godde et al. 2018). This method involves calculating the luminance difference between two patches of an image and dividing it by the total luminance of both patches. Luminance values were determined using a custom Matlab script, which averaged pixel intensity values from specified coordinates for all black and white stripes.

#### Statistical Analysis

We tested if turbidity source and level had a statistically significant effect on luminance, hue, chroma, image contrast, pH, and KH. We also tested whether turbidity source had a significant effect on turbidity level over time, which could indicate whether some types of turbidity remain suspended more than others. In total, we conducted seven different statistical tests (see Table 1 for model parameters). All statistical analyses were conducted using R Studio (v.4.3.2) using the lme4 (linear mixed models) package (Bates 2015). To identify models that best fit our data, we tested various model parameters and interactions and compared the resulting models using Akaike Information Criterion (AIC). We selected the model with the lowest AIC value and then validated it using the DHARMa package, which tests for residual normality, over-dispersion, and outliers (Hartig and Lohse 2022). Some models did not meet the assumption of residual normality (see table 1), but each model was selected as the best possible fit model amongst those tested (Hartig and Lohse 2022) (see Supplementary S3 for model selection). Results were plotted using the ggplot2 package (Wickham 2016).

**Table 1:**
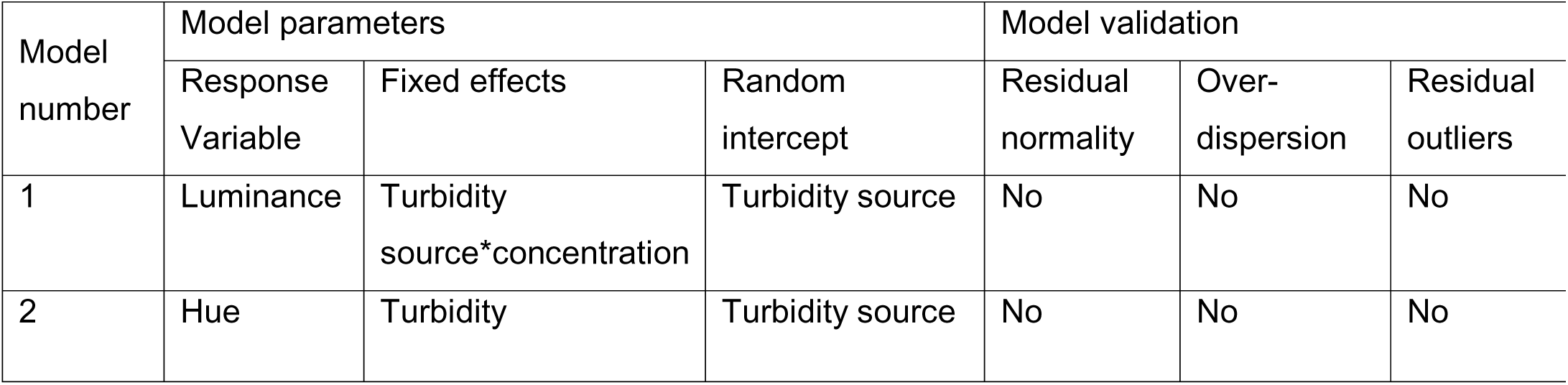
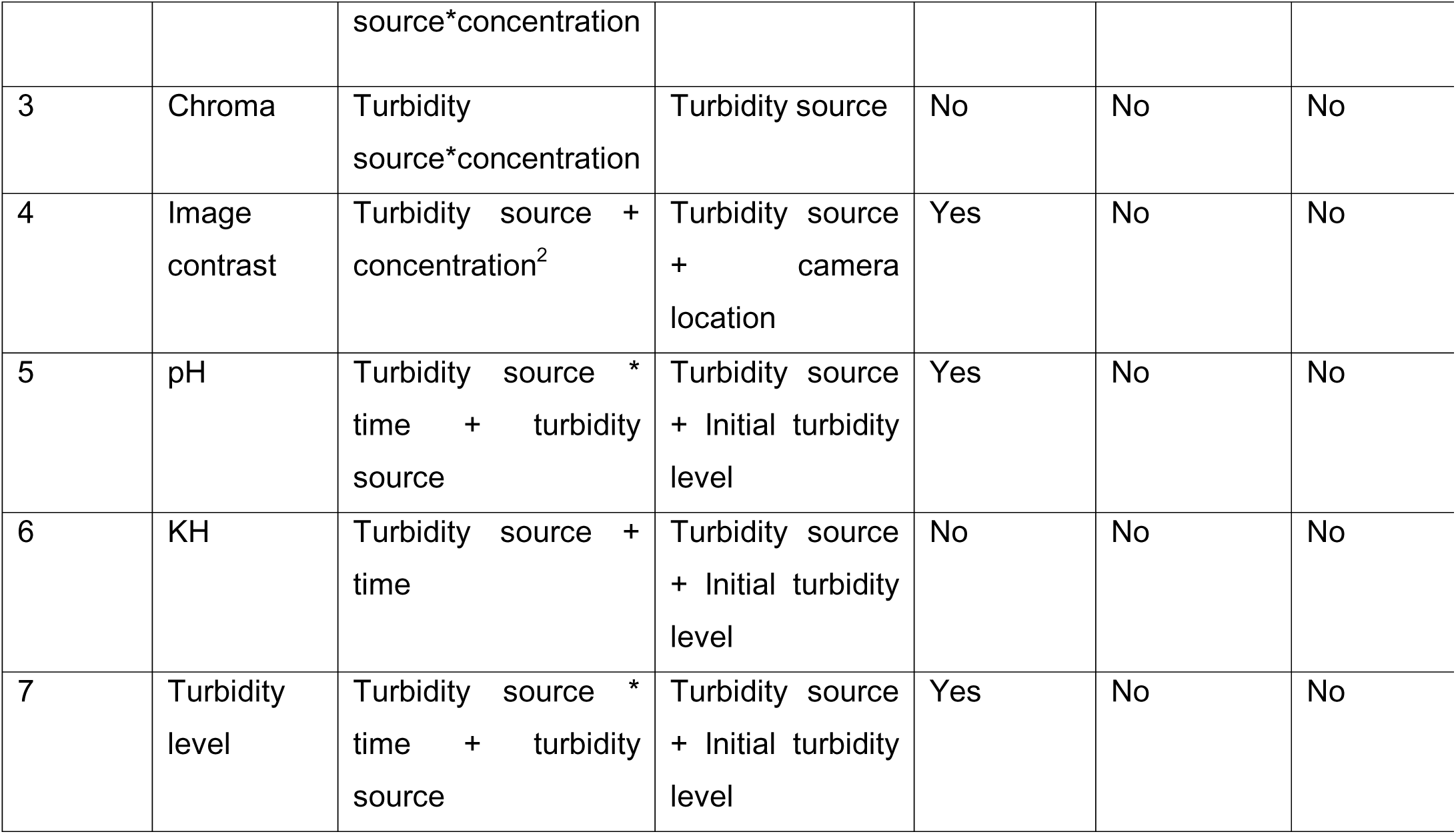
Summary of linear model parameters. For well-fitting models, residual normality is expected (‘Yes’), no overdispersion (‘No’), and no residual outliers (‘No’).

## Results

### Effect of turbidity on absolute irradiance

Changes in the shape of a spectral curve can provide insights into the light environment. Fig. 2A-D shows that turbidity source and level can alter the total irradiance (i.e. total number of photons). Increasing turbidity level generally increases the number of photons for bentonite, calcium carbonate, and kaolin, whereas algae have the opposite effect. Changes to the shape of the curve can provide information on the number of photons at different wavelengths and indicate an overall change in the colour of the light environment. These changes in shape can be seen in Fig 2A-D but are more obvious in Fig 2E-H where total irradiance has been standardized. In the presence of algae, there are fewer photons in the short-wavelength region (∼400-525 nm) and more photons in the medium-to-long wavelengths (∼525-650 nm). Bentonite similarly reduced the number of photons at shorter wavelengths while increasing them at longer wavelengths (∼575-700 nm), with a slight shift toward longer wavelengths.

Turbidity level has an effect for bentonite, as the shape of the curve between ∼575-700 nm, differs from 10 NTU to 94 NTU. Algae also shows an effect with turbidity, as the curve shape between ∼400-525 nm and ∼525-650 nm differs between 10 NTU and 52 NTU. In contrast, calcium carbonate and kaolin show minimal change from the clear water condition, with only slight changes at the highest turbidity levels.

### Effect of turbidity on luminance, hue, and chroma

We found a significant, non-linear relationship between luminance and turbidity level (Supplementary S4, Table 2). There was a significant interaction between all turbidity sources and turbidity level on luminance: increasing turbidity level caused an increase in luminance for kaolin (8.02×10^18^ photons ± SE = 8.64 × 10^17^, p = <0.001), followed by calcium carbonate (7.15 × 10^18^ photons ± SE = 8.00 × 10^17^, p = <0.001). Bentonite resulted in a smaller but still significant increase (3.77×10^18^ photons ± SE = 7.97 × 10^17^, p = <0.001), while algae led to a significant decrease in luminance (−1.99 × 10^18^ photons ± SE = 8.13 × 10^17^, p = 1.47 × 10^−2^) (F**ig. 2I**).

There was a significant interaction between all turbidity sources and turbidity level on hue: increasing turbidity level caused the greatest shift in hue for bentonite (6.31×10^−3^° ± SE = 3.35 × 10^−4^, p= <0.001), followed by algae (5.40 × 10^−3^° ± SE = 8.53 × 10^−4^, p= <0.001), then calcium carbonate (1.96 × 10^−3^ ° ± SE = 3.99 × 10^−4^, p= <0.001), then kaolin (1.51 × 10^−3^° ± SE = 4.11 × 10^−4^, p= <0.001) (**Fig. 2J**, Supplementary S4, Table 3).

Furthermore, there was a significant interaction between some turbidity sources and their level on chroma: increasing turbidity level caused an increase in chroma for algae (3.32 × 10^−3^ ± SE = 2.68 × 10^−4^, p= <0.001) and bentonite (3.71 × 10^−4^ ± SE = 1.05 × 10^−4^, p = 1.39 × 10^−3^). Neither calcium carbonate (−1.34 × 10^−4^ ± SE = 1.25 × 10^−4^, p = 0.29) nor kaolin (−1.68 × 10^−4^ ± SE = 1.29 × 10^−4^, p = 0.20) turbidity level had a significant impact on chroma (**Fig. 2K**, Supplementary S4, Table 4).

We found a significant, relationship between image contrast (Michelson contrast) and turbidity level (6.86 × 10^−3^ ±SE = 7.84 × 10^−4^, p <0.001) (**Fig. 2L**, Supplementary S4, Table 5). When considering individual turbidity sources, algae had the greatest effect on image contrast (−0.69 ± SE = 6.06 × 10^−2^, p <0.001), followed by kaolin (−0.59 ± SE = 5.97 × 10^−2^, p <0.001), then bentonite (−0.46 ± SE = 5.88 × 10^−2^, p <0.001), and calcium carbonate (−0.37 ± SE = 5.96 × 10^−2^, p <0.001).

### Turbidity settling rate, pH and water hardness over time

The settling rate of each turbidity was described by repeatedly measuring suspended turbidity over time. All turbidity led to some degree of settling, but there was a significant effect of turbidity source on the settling rate (**Fig. 2M**, Supplementary S4, Table 6). Calcium carbonate had the most rapid settling rate (−0.24 NTU ± SE = 2.73 × 10^−2,^ p <0.001), followed by algae (− 0.15 NTU ± SE = 3.79 × 10^−2^, p <0.001), kaolin (−8.21 NTU × 10^−2^ ± SE = 1.73 × 10^−2^, p <0.001), and bentonite (−6.47 × 10^−2^ NTU ± SE = 1.73 × 10^−2^, p <0.001).

Some turbidity sources had a significant effect on pH, as well as a significant interaction between turbidity source and time. Compared to pure RO water (pH=6.73 ± SE = 0.22), both algae and calcium carbonate significantly increased pH (1.91 ± SE = 0.37, p < 0.001 and 1.33 ± SE = 0.32, p < 0.001, respectively) but bentonite and kaolin had non-significant effects (p > 0.05) (**Fig. 2N**, Supplementary S4, Table 7). Over time, calcium carbonate significantly increased pH (0.19 ±SE = 4.60 × 10^−2^, p < 0.001), while kaolin decreased pH (0.12 ±SE = 2.91 × 10^−2^, p < 0.001). The observed trend was that changes in pH generally occurred within the first hour, except in the case of algae (**Fig. 2N**).

There was a significant impact of some turbidity sources on the water KH. KH increased for calcium carbonate (1.50 ±SE = 0.19, p < 0.001) and algae (1.00 ±SE = 0.23, p < 0.001), but not bentonite or kaolin (p>0.05) **(Fig. 2O**, Supplementary S4, Table 8).

## Discussion

As land use changes and global temperatures rise, turbidity conditions in both freshwater and marine habitats are expected to shift, impacting not only turbidity levels but also their sources. As a result, fish will increasingly encounter turbidity levels and sources beyond what they have evolved to cope with. Our results demonstrate that turbidity’s effects on the light environment are significant and dependent on both turbidity level and source. Algae reduce luminance, shift hue towards medium-long wavelengths, and increase chroma, creating a darker, greener, and more saturated light environment. Bentonite slightly increases luminance, shifts hue towards medium-long and long wavelengths, and slightly increases chroma, resulting in a brighter, redder light environment. Calcium carbonate and kaolin increase luminance but have minimal effects on hue and chroma, producing a much brighter environment without altering chromatic properties. These findings have implications for assessing the risks of changing turbidity to natural environments. For example, Engström-Öst and Candolin (2007) found that increasing algae-based turbidity can alter courtship behaviour in three-spined sticklebacks, and males in turbid water had to work harder for female attention. As part of courtship, males of this species develop a bright red belly during the breeding season, which serves as an important fitness signal (Seehausen, Terai et al. 2008). Our results here predict that the red patches would be less conspicuous in algal turbidity but remain visible in bentonite-induced turbidity. Thus, for this species, nutrient runoff leading to algal blooms may impair courtship behaviours, while clay soil runoff is less likely to have such an impact. This study highlights the need to consider turbidity’s impact on the visual environment when trying to understand and predict its effects on aquatic species and their behaviour.

Although turbidity is generally associated with reduced light levels, our experiments show that the total light available was actually higher in turbid water for the inorganic turbidity sources (Bowmaker 1995) tested. Both irradiance (Fig 2A-D) and luminance (Fig 2I) were greater in turbid water than in clear water. This increase in brightness is due to light scattering, which is influenced by factors such as particle composition, size, concentration (Utne-Palm 2002, Johnsen 2012), as well as other physical water properties including temperature, salinity, and pressure (Zhang and Hu 2021, Tomperi, Isokangas et al. 2022). In contrast, algal turbidity reduces light availability, as photosynthetic cells absorb rather than scatter light. The achromatic component of the light environment (i.e. luminance) guides many fundamental visual tasks such motion detection, depth perception, and pattern and texture discrimination (Siebeck, Wallis et al. 2014, van den Berg, Hollenkamp et al. 2020). A shift in luminance could prevent the detection of critical visual cues. Furthermore, a significant decrease in luminance may cause a shift from cone-based to rod-based vision (Wolfe and Ali 2015), which would substantially impact visual abilities. As rods generally have lower acuity than cones (Bowmaker 1995), significantly reduced luminance could impair a species’ ability to detect smaller visual cues or fine features. This is likely to happen with some turbidity sources, such as charcoal, that lead to a rapid drop in luminance (e.g. by 1190% at 13.3 NTU, unpublished data).

Light consists of both achromatic (luminance) and chromatic (hue, chroma) components. Spectra with a decrease in the area under the curve will have a lower luminance, making the environment appear darker. The luminance of a visual feature is proportional to how closely its reflectance spectrum matches the ambient light spectrum. For this reason, a blue fish in blue water will appear brighter (higher luminance) and more visible than a red fish. Changes to the chromatic component of the light environment shift the relative availability of different wavelengths, altering how visual stimuli reflecting those wavelengths appear (Endler 1990). Spectra with differing wavelengths of greatest slope are perceived as different hues (Endler 1990), and spectra with a greater difference between the maximum and minimum peaks of the curve will have a greater chroma, resulting in more saturated colours. Additionally, the perceived hue of a visual feature can change based on the spectral properties of the turbidity source, with these effects being wavelength-specific. Turbidity sources like algae, which absorb longer wavelengths (e.g. red, orange), reduce the visibility of red and orange features, while green and blue features will remain conspicuous. In contrast, bentonite turbidity scatters shorter wavelengths (e.g. blue, green), limiting the visibility of blue and green features but not red, orange or brown ones. Both algae and bentonite also significantly increase the chroma of the light environment, so that the light environment is more saturated in specific regions of the spectrum. The saturated region is dependent on the turbidity source. Regardless of a species’ visual system, spectra with a greater magnitude of change between different parts of the spectrum will typically be perceived as having greater chroma, and spectra with differing wavelengths of greatest slope will be perceived as different hues (Endler 1990). Finally, we observed a significant decrease in image contrast with the addition of all turbidity sources and increasing turbidity levels. Image contrast measurements incorporate estimates from all four optical factors (i.e. the ambient light environment, the spectral reflectance of the visual stimuli, the transmission of light through the medium, and the amount of veiling light) in the light pathway to provide an approximation of how turbidity affects the visibility of a stimulus and therefore can serve as a practical tool for assessing the impacts of turbidity without complex visual modelling. In our setup, the greatest changes in image contrast, and consequently stimulus visibility, occurred between 0-15 NTU, with effects stabilizing at 20 NTU. The combined chromatic changes to the light environment are likely to significantly affect visually-guided behaviours in fish, particularly those relying on colour signals such as sexual selection, aposematism, spatial cognition, camouflage, and predator-prey interactions (Endler 1993).

While we suggest that experimental turbidity sources should be selected for their potential effects on the visual environment, not all sources will be suitable for all species. When turbidity sources were added to RO water, we observed a significant drop in turbidity level over time that was independent of the turbidity source. Calcium carbonate had the greatest turbidity level change over time, followed by algae, kaolin, then bentonite. The difference in turbidity level change over time between turbidity sources found here is useful to consider when designing turbidity experiments. Scientists wishing to take advantage of quickly changing turbidity levels, for example where changing turbidity levels between trials is logistically challenging, might choose to use a turbidity source that settles quickly. Turbidity levels that change more slowly on the other hand, might be more useful to experiments with longer trial durations. Both pH and KH were significantly impacted in a turbidity source-dependent manner. Algae, calcium carbonate and kaolin lead to significant increases or decreases (for kaolin only) in the water pH. While the change in pH was immediate with algae, it was observed as soon as the second measurement was taken (one hour after the turbidity source was added). This indicates that these calcium carbonate and kaolin need a short time in the water to modify its chemistry. Calcium carbonate and algae significantly increased the water KH. The results of this study are not intended to allow researchers to predict the precise impact of each turbidity source on pH and KH in their experimental design, as small changes in their design such as water mineral content and temperature can impact pH and KH (Irwin, Stoner et al. 2013). However, it demonstrates that pH and KH must be taken into account when designing turbidity experiments and monitored during experiments as they may change with turbidity source and time and could crucially impact the welfare of the study species and the output of behavioural experiments (Kleinhappel, Burman et al. 2019, Cleal, Gibbon et al. 2020). This paper is intended to serve as a guide to key considerations and potential effect sizes, acknowledging that each environment, natural or laboratory, has unique characteristics.

This study explores how turbidity influences the light environment, and demonstrates that luminance, hue, chroma, and image contrast are significantly affected by turbidity, with variation depending on turbidity source and level. Our findings indicate that different sources and levels of turbidity cannot be assumed to uniformly affect the appearance of visual stimuli or the aquatic environment. We also examined particle setting rate by turbidity source, along with their effects on pH and KH. Understanding turbidity’s impact on animals requires considering how it alters visual information and may help us identify the mechanisms species have evolved to cope with changing visual conditions. Moreover, this study highlights the need for conservation ecology studies to consider turbidity levels, sources, and ecological contexts when interpreting the potential consequences of turbidity for wild fish populations.

## Acknowledgements

We thank animal technicians and field station support staff: Helen Sanders, Christine Soper, Allex Turner, Phil Smith, John Hogg, and Angela Bishop.

## Data and resource availability

We are creating a data repository link at the time of submission linked to the manuscript. In the meantime, all data and code are available on demand.

