## Supplementary Information for "A sensory approach to turbidity: How sources and levels shape aquatic light environments and fish visual ecology"

### S1 Review:

A literature review was conducted to assess the prevalence of the use of different turbidity sources and fish response categories in studies that explore how fish respond to turbidity. The search was conducted using the Web of Science database using the following search string for papers with titles that match: (“turbidity” OR “suspended sediment”) AND (“fish” OR “teleost” OR “behav*” OR “predat*” OR “antipredat*” OR “prey” OR “reactive distance” OR “optomotor” OR “fitness” OR “surviv*” OR “courtship” OR “reproduct*” OR “collective” OR “avoidance” OR “forag*” OR “move*” OR “navigat*” OR “swim*” OR “len*” OR “metabol*” OR “gill” OR “diversity” OR “sex*”). From this search, 398 references were found, and a further 22 references were identified by reviewing the cited literature in papers published within the last year from the initial string search. 200 references were included, with others removed as they either did not include measurements of fish or turbidity, or were only theoretical/ review articles, or were unreleased papers, or were for conference abstracts for a talk, or were a PhD thesis subsequently published as separate papers. Papers were organised by reference, category of response variable, and turbidity source used.

Table 1: Number of studies from review per field per turbidity source.

| | algae/ phytoplankton | bentonite | clay | kaolin | natural sediment | calcium carbonate | diatomite / silica | palm oil mill effluent |
|---|---|---|---|---|---|---|---|---|
| Antipredator behaviour | 3 | 7 | 2 | 6 | 9 | 0 | 0 | 0 |
| Biodiversity / Distribution | 0 | 0 | 1 | 0 | 14 | 0 | 0 | 0 |
| Foraging efficiency | 13 | 26 | 17 | 11 | 23 | 1 | 3 | 1 |
| Movement behaviour | 1 | 4 | 2 | 1 | 5 | 1 | 3 | 0 |
| Physiological condition | 4 | 4 | 2 | 1 | 12 | 1 | 0 | 0 |
| Sexual selection | 15 | 3 | 0 | 0 | 1 | 0 | 0 | 0 |
| Social / Collective behaviour | 0 | 3 | 0 | 7 | 3 | 0 | 0 | 0 |

### S2 Experimental setup:

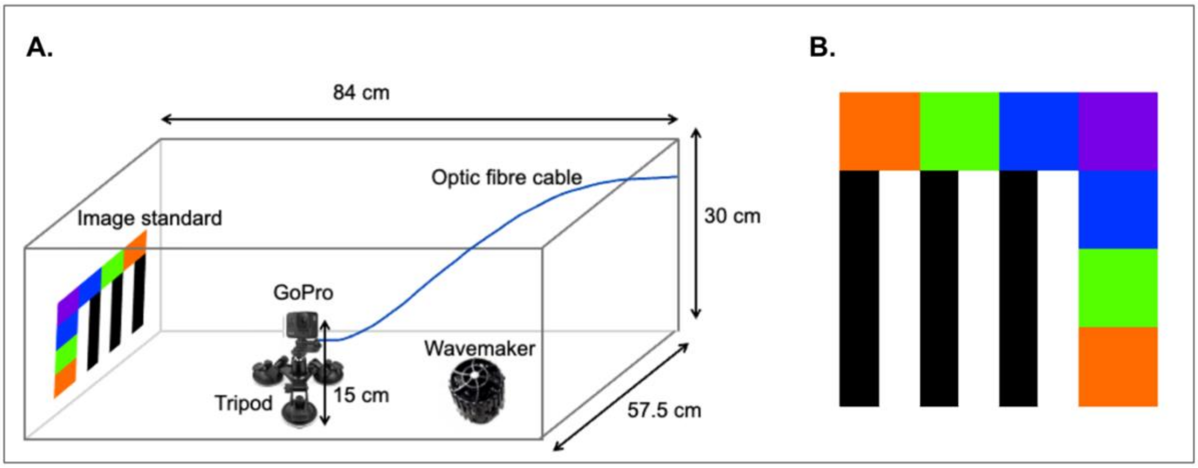


**Figure 1 - Experimental Setup: (A)** Experimental Tank (84 cm x 30 cm x 57.5 cm; l x h x w) filled with 135 litres of water with a tripod placed in its centre and the GoPro facing an image standard on the far wall. An optic fibre cable (blue) was secured to the camera with a custom printed holder and is pointed towards the image standard. **(B)** Image Standard: A laminated printed card (30 x 30 cm) containing three pairs of horizontal black and white gratings, with a row of different coloured squares above, and a column of the coloured squares to the left of the gratings. The colours include two that the cones of goldfish would be maximally sensitive to (Hex colour: #0035ff, #ff6b00), and two that triggerfish cones would be maximally sensitive to (Hex colour: #7900e6, #56ff00).

### S3 Model Selection:

Luminance: The best fitting model included the fixed effects of an interaction between turbidity source and turbidity level, and the second order polynomial of turbidity level, and a random effect of turbidity source. Transforming the luminance data did not improve the model validity. From all models tested, no models fulfilled the assumption of normality of residuals, but the selected model fit the assumptions of homoscedasticity and no outliers of residuals. As this was the most valid model, AIC scores were not compared between other models.

Hue: The best fitting model included the fixed effects of an interaction between turbidity source and turbidity level, and a random effect of turbidity source. Transforming the hue data did not improve the model validity. From all models tested, no models fulfilled the assumption of normality of residuals, but the selected model successfully fulfilled the assumptions of homoscedasticity and no outliers in its residuals. As transformations to the response variable did not improve the model validity, only the models with non-transformed data were considered. The first two models of the model set did not have sufficient sample size to include all of the fixed effects and interactions. The remaining two models were compared in their AIC values using an ANOVA test and the model with the smaller value chosen.

Chroma: Similarly to hue, the best fitting model included the fixed effects of an interaction between turbidity source and turbidity level, and a random effect of turbidity source. Transforming the chroma data did not improve the model validity. From all models tested, no models fulfilled the assumption of normality of residuals, but the selected model fit the assumptions of homoscedasticity and no outliers of residuals. As transformations to the response variable did not improve the model validity, only the models with non-transformed data were considered. The first two models of the model set did not have sufficient sample size to include all of the fixed effects and interactions. The remaining two models were compared in their AIC values using an ANOVA test and the model with the smaller value chosen.

Image Contrast: The best fitting model included the fixed effects of turbidity level, turbidity source, and the second order polynomial term of turbidity level, and random effects of turbidity source and camera location. Performing transformations on the image contrast variable did not improve the model validity. The chosen model fulfilled all assumptions of normality of residuals, homoscedasticity, and no outliers of residuals confirmed using the DHARMa package. This model also displayed the lowest AIC from those models with similar model validity.

Turbidity level: The best fitting model included the fixed effects of an interaction between turbidity source and time, and turbidity source alone, and random effects of turbidity source and starting turbidity level. The model that best met the model assumptions included a log transformation applied to the turbidity level response variable. From all models tested, three models with a log transformation, and three models with a square root transformation fit all model assumptions. Visual inspection of the histograms of the transformations of the response variable was used to select the log transformed model set for further assessment. The final selected model had the lowest AIC score of the model set, and had the fewest number of terms.

pH: The best fitting model included the fixed effects of an interaction between turbidity source and time, and turbidity source alone, and random effects of turbidity source and starting turbidity level. Transforming the response variable did not improve the model validity. From all models tested, only one model fulfilled all model assumptions confirmed using the DHARMa package. The final selected model had the lowest AIC score of the model set, and had the fewest number of terms.

KH: The best fitting model included the fixed effects of turbidity source and time, and random effects of turbidity source and starting turbidity level. Transforming the response variable did not improve the model validity. From all models tested, none fit any of the model assumptions of normality of residuals, but the selected model fit the final two assumptions of homoscedasticity and no significant outliers of residuals. The first two models of the model set did not have sufficient sample size to include all of the fixed effects and interactions. From the remaining models in the model set, the model with the lowest AIC value was chosen.

### S4 Model Outputs:

Table 2 - Output of best fitting model of luminance. Results from linear mixed effects model. Significant results are in italics. SE = standard error, df = degrees of freedom, s.d. = standard deviation. n = 37. Intercept corresponds to a turbidity level of 0 NTU for the no turbidity (clear water) condition.

**Luminance**

| Fixed Effects | Estimate | SE | df | t value | p-value |
|---|---|---|---|---|---|
| (Intercept) | 5.68E+20 | 8.13E+19 | 3.05 | 6.98 | 5.71E-03 |
| I(Turbidity Level^2) | -4.79E+16 | 7.65E+15 | 3.08E+05 | -6.26 | 3.76E-10 |
| No Turbidity : Turbidity Level | -2.47E+19 | 8.26E+19 | 3.09 | -0.30 | 0.78 |
| Algae : Turbidity Level | -1.99E+18 | 8.13E+17 | 3.05E+05 | -2.44 | 1.47E-02 |
| Bentonite : Turbidity Level | 3.77E+18 | 7.97E+17 | 2.34E+05 | 4.73 | 2.26E-06 |
| Calcium Carbonate : Turbidity Level | 7.15E+18 | 8.00E+17 | 1.66E+05 | 8.94 | < 2e-16 |
| Kaolin : Turbidity Level | 8.02E+18 | 8.64E+17 | 6.94E+04 | 9.28 | < 2e-16 |
| **Random Effects** | **Name** | **Variance** | **s.d.** | | |
| Turbidity Source | (Intercept) | 2.58E+40 | 1.61E+20 | | |
| Residual | | 9.27E+38 | 3.05E+19 | | |

Table 3 - Output of best fitting model of hue. Results from linear mixed effects model. Significant results are in italics. SE = standard error, df = degrees of freedom, s.d. = standard deviation. n = 37. Intercept corresponds to a turbidity level of 0 NTU for the no turbidity (clear water) condition.

**Hue**

| Fixed Effects | Estimate | SE | df | t value | p-value |
|---|---|---|---|---|---|
| (Intercept) | 0.70 | 8.18E-02 | 2.98 | 8.55 | 3.45E-03 |
| No Turbidity : Turbidity Level | -4.20E-02 | 8.37E-02 | 3.13 | -0.50 | 0.65 |
| Algae : Turbidity Level | 5.40E-03 | 8.53E-04 | 29.59 | 6.33 | 6.01E-07 |
| Bentonite : Turbidity Level | 6.31E-03 | 3.35E-04 | 29.37 | 18.87 | < 2e-16 |

| | | | | | |
|---|---|---|---|---|---|
| Calcium Carbonate : Turbidity Level | 1.96E-03 | 3.99E-04 | 29.46 | 4.91 | 3.11E-05 |
| Kaolin : Turbidity Level | 1.51E-03 | 4.11E-04 | 29.64 | 3.69 | 9.12E-04 |
| Random Effects | Name | Variance | s.d. | | |
| Turbidity Source | (Intercept) | 2.63E-02 | 0.16203 | | |
| Residual | | 1.30E-03 | 3.61E-02 | | |

Table 4 - Output of best fitting model of chroma. Results from linear mixed effects model. Significant results are in italics. SE = standard error, df = degrees of freedom, s.d. = standard deviation. n = 37. Intercept corresponds to a turbidity level of 0 NTU for the no turbidity (clear water) condition.

**Chroma**

| Fixed Effects | Estimate | SE | df | t value | p-value |
|---|---|---|---|---|---|
| (Intercept) | 0.30 | 2.57E-02 | 2.98 | 11.60 | 1.41E-03 |
| No Turbidity : Turbidity Level | -9.83E-03 | 2.63E-02 | 3.13 | -0.37 | 0.73 |
| Algae : Turbidity Level | 3.32E-03 | 2.68E-04 | 29.59 | 12.37 | 3.22E-13 |
| Bentonite : Turbidity Level | 3.71E-04 | 1.05E-04 | 29.38 | 3.53 | 1.39E-03 |
| Calcium Carbonate : Turbidity Level | -1.34E-04 | 1.25E-04 | 29.46 | -1.07 | 0.29 |
| Kaolin : Turbidity Level | -1.68E-04 | 1.29E-04 | 29.65 | -1.30 | 0.20 |
| Random Effects | Name | Variance | s.d. | | |
| Turbidity Source | (Intercept) | 2.58E-03 | 0.05084 | | |
| Residual | | 1.28E-04 | 0.01133 | | |

Table 5 - Output of best fitting model of image contrast. Results from linear mixed effects model. Significant results are in italics. SE = standard error, df = degrees of freedom, s.d. = standard deviation. n = 103. Intercept corresponds to a turbidity level of 0 NTU for the no turbidity (clear water) condition.

**Image contrast**

| Fixed Effects | Estimate | SE | df | t value | p-value |
|---|---|---|---|---|---|
| (Intercept) | 0.78 | 6.75E-02 | 11.99 | 11.63 | 6.87E-08 |
| Turbidity Level | -6.86E-03 | 7.84E-04 | 94.41 | -8.75 | 7.96E-14 |
| Algae | -0.69 | 6.06E-02 | 92.03 | -11.31 | < 2e-16 |
| Bentonite | -0.46 | 5.88E-02 | 91.98 | -7.82 | 8.40E-12 |
| Calcium Carbonate | -0.37 | 5.96E-02 | 92.02 | -6.15 | 2.00E-08 |
| Kaolin | -0.59 | 5.97E-02 | 92.17 | -9.94 | 2.97E-16 |
| I(Turbidity Level^2) | 4.12E-05 | 6.77E-06 | 93.72 | 6.09 | 2.50E-08 |
| Random Effects | Name | Variance | s.d. | | |
| Camera Location | (Intercept) | 1.60E-03 | 3.99E-02 | | |
| Turbidity Source | (Intercept) | 1.25E-02 | 0.11 | | |
| Residual | | 1.91E-03 | 4.37E-02 | | |

Table 6 - Output of best fitting model of settling rate. Results from linear mixed effects model. Significant results are in italics. SE = standard error, df = degrees of freedom, s.d. = standard deviation. n = 105. Intercept corresponds to the no turbidity (clear water) condition at time = 0.

**Settling rate**

| Fixed Effects | Estimate | SE | df | t value | p-value |
|---|---|---|---|---|---|
| (Intercept) | 0.88 | 0.39 | 71.62 | 2.26 | 2.70E-02 |
| Algae | 2.06 | 0.66 | 43.24 | 3.10 | 3.40E-03 |
| Bentonite | 2.47 | 0.55 | 61.19 | 4.49 | 3.21E-05 |
| Calcium Carbonate | 2.72 | 0.67 | 43.74 | 4.08 | 1.90E-04 |
| Kaolin | 2.73 | 0.55 | 61.19 | 4.96 | 5.94E-06 |

| | | | | | |
|---|---|---|---|---|---|
| Time : No Turbidity | -4.02E-03 | 8.03E-02 | 47.48 | -0.05 | 0.96 |
| Time: Algae | -0.15 | 3.79E-02 | 84.24 | -4.07 | 1.06E-04 |
| Time : Bentonite | -6.47E-02 | 1.73E-02 | 82.29 | -3.75 | 3.32E-04 |
| Time : Calcium Carbonate | -0.24 | 2.73E-02 | 82.29 | -8.94 | 9.35E-14 |
| Time : Kaolin | -8.21E-02 | 1.73E-02 | 82.29 | -4.75 | 8.46E-06 |
| Random Effects | Name | Variance | s.d. | | |
| Starting Turbidity level | (Intercept) | 0.45 | 0.6731 | | |
| Turbidity Source | (Intercept) | 5.75E-02 | 0.2398 | | |
| Residual | | 4.18E-02 | 0.2044 | | |

Table 7 - Output of best fitting model of pH. Results from linear mixed effects model. Significant results are in italics. SE = standard error, df = degrees of freedom, s.d. = standard deviation. n = 105. Intercept corresponds to the no turbidity (clear water) condition at time = 0.

**pH**

| Fixed Effects | Estimate | SE | df | t value | p-value |
|---|---|---|---|---|---|
| (Intercept) | 6.73 | 0.22 | 94.50 | 30.51 | < 2e-16 |
| Algae | 1.91 | 0.37 | 78.27 | 5.16 | 1.85E-06 |
| Bentonite | -0.45 | 0.28 | 93.37 | -1.59 | 0.12 |
| Calcium Carbonate | 1.33 | 0.32 | 87.21 | 4.17 | 7.19E-05 |
| Kaolin | -0.38 | 0.28 | 93.37 | -1.35 | 0.18 |
| Time : No Turbidity | -4.26E-03 | 4.89E-02 | 86.25 | -8.70E-02 | 0.93 |
| Time: Algae | -0.11 | 0.06 | 84.88 | -1.70 | 9.22E-02 |
| Time : Bentonite | -5.08E-02 | 2.91E-02 | 84.88 | -1.75 | 8.42E-02 |
| Time : Calcium Carbonate | 0.19 | 0.05 | 84.88 | 4.08 | 1.00E-04 |
| Time : Kaolin | -0.12 | 0.03 | 84.88 | -4.07 | 1.04E-04 |
| Random Effects | Name | Variance | s.d. | | |
| Starting Turbidity level | (Intercept) | 1.57E-02 | 0.13 | | |
| Turbidity Source | (Intercept) | 1.77E-02 | 0.13 | | |

| | | | |
|---|---|---|---|
| Residual | | 0.12 | 0.34 |

Table 8 - Output of best fitting model of KH. Results from linear mixed effects model. Significant results are in italics. SE = standard error, df = degrees of freedom, s.d. = standard deviation. n = 105. Intercept corresponds to the no turbidity (clear water) condition at time = 0.

**KH**

| Fixed Effects | Estimate | SE | df | t value | p-value |
|---|---|---|---|---|---|
| (Intercept) | 1.02 | 0.10 | 94.24 | 9.79 | 4.97E-16 |
| Algae | 1.00 | 0.23 | 33.85 | 4.29 | 1.39E-04 |
| Bentonite | 8.62E-02 | 0.16 | 71.48 | 0.55 | 0.58 |
| Calcium Carbonate | 1.50 | 0.19 | 46.61 | 7.96 | 3.06E-10 |
| Kaolin | 4.53E-04 | 0.16 | 71.48 | 3.00E-03 | 1.00 |
| Time | -7.77E-03 | 7.32E-03 | 88.85 | -1.06 | 0.29 |
| **Random Effects** | **Name** | **Variance** | **s.d.** | | |
| Starting Turbidity level | (Intercept) | 3.46E-02 | 0.19 | | |
| Turbidity Source | (Intercept) | 6.27E-03 | 7.92E-02 | | |
| Residual | | 2.07E-02 | 0.14 | | |

### S5 Code:

```
library(readr)
library(dplyr)
library(ggplot2)
library(viridis)
library(gridExtra)
library(grid)
library(ggh4x)
library(scales)
library(reshape2)
library(purrr)
library(tidyr)
library(pracma)
library(DHARMa)
library(lme4)
library(lmerTest)
library(report)


#-------------------------------------#
#-----------   Load Data    -----------#
#-------------------------------------#
old_cal_file <- file.choose() #choose old_cal file
new_cal_file <- file.choose() #choose new_cal file
Turbidity_source_folder_path <- dirname(file.choose()) #choose a file in Turbidity source data
folder
Algae_folder_path <- dirname(file.choose()) #choose a file in Algae folder
Turbidity_Level_data<-read.csv(file.choose()) %>% #choose Turbidity_Level_data
  dplyr::select(Turbidity_Source, Measurement, Average_NTU)
Turbidity_Level_data$Average_NTU <- as.numeric(Turbidity_Level_data$Average_NTU)

Exp_design<-read.csv(file.choose()) %>% #choose Experimental_design_variables file
  dplyr::select(Camera_Location, Time, Turbidity_Source, Measurement, Average_NTU,
Starting_Turbidity_level, pH, KH, Michelson_Contrast, Date)
Exp_design$Turbidity_Source<-as.factor(Exp_design$Turbidity_Source)
```

```
Exp_design$Camera_Location<-as.factor(Exp_design$Camera_Location)
Exp_design$Starting_Turbidity_level<-as.factor(Exp_design$Starting_Turbidity_level)
Exp_design$Date<-as.factor(Exp_design$Date)
baseline_avgpH <- mean(Exp_design$pH[Exp_design$Turbidity_Source == "Baseline"]) #
Calculate the average NTU for the "Baseline" group
baseline_avgKH <- mean(Exp_design$KH[Exp_design$Turbidity_Source == "Baseline"]) #
Calculate the average NTU for the "Baseline" group
baseline_avgIC <- mean(Exp_design$Michelson_Contrast[Exp_design$Turbidity_Source ==
"Baseline"]) # Calculate the average NTU for the "Baseline" group
baseline_avgSR <- mean(Exp_design$Average_NTU[Exp_design$Turbidity_Source ==
"Baseline"]) # Calculate the average NTU for the "Baseline" group
Exp_design_1 <- Exp_design %>% filter(Turbidity_Source != 'Baseline')


##########################################################################
########################################################################
#--------------------------------------------------------------------------------------------------------------
---------------------#
#                                        Remove old calibration, add new calibration
#--------------------------------------------------------------------------------------------------------------
---------------------#
##########################################################################
########################################################################
#Original calibration of absolute irradiance measurements was incorrect so this was removed
and the correct calibration applied

#-----------------------------------#
#------  Perform Recalibration    -----#
#-----------------------------------#
#Old Cal Turbidity Sources: Bentonite, Calcium.Carbonate, Charcoal, and Kaolin
#Function to process each file
process_file <- function(file_path, old_cal, new_cal) {
  # Read the file
  data <- read_csv(file_path)
  # Remove rows where Wavelength values are less than 400 or more than 700
  data <- data %>%
    filter(Wavelength >= 400 & Wavelength <= 700)
```

```
  # Merge with old calibration data
  data <- left_join(data, old_cal, by = "Wavelength")
  # Merge with new calibration data
  data <- left_join(data, new_cal, by = "Wavelength")
  # Calculate raw_values
  data$raw_values <- data$Absolute_Irradiance / data$old_cal
  # Calculate new_cal_values
  data$new_cal_values <- data$raw_values * data$new_cal
  # Calculate photons
  data$photons <- data$new_cal_values * data$Wavelength * 5.05e15
  # Determine Turbidity_Source based on file name
  turbidity_source <- ifelse(grepl("Bentonite", file_path), "Bentonite",
                             ifelse(grepl("Calcium.Carbonate", file_path), "Calcium.Carbonate",
                                    ifelse(grepl("Kaolin", file_path), "Kaolin",
                                           ifelse(grepl("Charcoal", file_path), "Charcoal",
                                                  ifelse(grepl("No_Turbidity", file_path), "No_Turbidity", NA)))))
  # Determine Measurement based on file name
  measurement <- gsub("[^0-9]", "", basename(file_path))
  # Create a table with the required columns
  result <- data.frame(
    file_name = basename(file_path),
    Wavelength = data$Wavelength,
    photons = data$photons,
    Turbidity_Source = turbidity_source,
    Measurement = as.integer(measurement))
  return(result)}

old_cal <- read_csv(old_cal_file)
new_cal <- read_csv(new_cal_file)

# Get file paths in the folder
file_paths <- list.files(Turbidity_source_folder_path, full.names = TRUE)
# Initialize a list to store results
results <- list()
# Process each file
for (i in seq_along(file_paths)) {
  file_path <- file_paths[i]
```

```
  result <- process_file(file_path, old_cal, new_cal)
  results[[i]] <- result}
# Combine results into a single data frame
all_spectra <- do.call(rbind, results)


#Algae data
#Function to process each file
process_file1 <- function(file_path) {
  # Read the file
  data <- read_csv(file_path)
  # Remove rows where Wavelength values are less than 400 or more than 700
  data <- data %>%
    filter(Wavelength >= 400 & Wavelength <= 700)
  # Calculate photons
  data$photons <- data$photons * data$Wavelength * 5.05e15
  # Determine Turbidity_Source based on file name
  turbidity_source <- ifelse(grepl("Algae", file_path), "Algae",
                             ifelse(grepl("No_Turbidity", basename(file_path)), "No_Turbidity", NA))
  # Determine Measurement based on file name
  measurement <- gsub("[^0-9]", "", basename(file_path))
  # Create a table with the required columns
  result <- data.frame(
    file_name = basename(file_path),
    Wavelength = data$Wavelength,
    photons = data$photons,
    Turbidity_Source = turbidity_source,
    Measurement = as.integer(measurement))
  return(result)}

# Get file paths in the folder
file_paths <- list.files(Algae_folder_path, full.names = TRUE)
# Initialize a list to store results
results <- list()
# Process each file
for (i in seq_along(file_paths)) {
  file_path <- file_paths[i]
```

```
  result <- process_file1(file_path)
  results[[i]] <- result}
# Combine results into a single data frame
algae_spectra <- do.call(rbind, results)
# Change Turbidity_Source to "No_Turbidity" if "No" is in file_name
algae_spectra$Turbidity_Source[grepl("No", algae_spectra$file_name)] <- "No_Turbidity"

# Merge algae_spectra with all_spectra based on common columns
all_spectra <- bind_rows(all_spectra, algae_spectra)

# Merge final_result with Turbidity_Level_data
all_spectra <- all_spectra %>%
  left_join(Turbidity_Level_data, by = c("Turbidity_Source", "Measurement"))
# Print final result
print(all_spectra)

#------------No Turbidity adjusted photons calculations----------#
no_turbidity_spectra <- all_spectra[all_spectra$Turbidity_Source == 'No_Turbidity',]
# Function to calculate the area under the curve for each Turbidity_Source
calculate_area <- function(data) {
  area <- trapz(data$Wavelength, data$photons)
  return(data.frame(Area_Under_Curve = area))}
area_under_curve_data <- no_turbidity_spectra %>%
  group_by(Measurement) %>%
  nest() %>%
  mutate(area_data = map(data, calculate_area)) %>%
  unnest(cols = c(area_data))
print(area_under_curve_data)

#makes the area under each curve equaled to 1
no_turbidity_spectra1 <- no_turbidity_spectra %>%
  mutate(standardized_photons = case_when(
    Measurement == '1' ~ photons /  6.29e19,
    Measurement == '2' ~ photons / 7.58e19,
    Measurement == '3' ~ photons / 6.18e19,
    Measurement == '4' ~ photons / 6.83e19,
    Measurement == '5' ~ photons / 1.75e20,
```

```
    TRUE ~ photons  # For other cases, keep the original value
  ))%>%
  group_by(Wavelength)

no_turbidity_spectra1$Measurement<-as.factor(no_turbidity_spectra$Measurement)
ggplot(no_turbidity_spectra1, aes(x = Wavelength, y = standardized_photons, color =
Measurement)) +
  geom_line() +
  labs(x = "Wavelength (nm)", y = "Standardized photons/cm^2/nm", color = "Turbidity Source")
+
  theme_classic()

no_turb_1<-no_turbidity_spectra %>%
  filter(Measurement=='1')%>%
  dplyr::select(Wavelength, photons)%>%
  rename(Measurement_1 = photons)
no_turb_2<-no_turbidity_spectra %>%
  filter(Measurement=='2')%>%
  dplyr::select(Wavelength, photons)%>%
  rename(Measurement_2 = photons)
no_turb_3<-no_turbidity_spectra %>%
  filter(Measurement=='3')%>%
  dplyr::select(Wavelength, photons)%>%
  rename(Measurement_3 = photons)
no_turb_4<-no_turbidity_spectra %>%
  filter(Measurement=='4')%>%
  dplyr::select(Wavelength, photons)%>%
  rename(Measurement_4 = photons)
no_turb_5<-no_turbidity_spectra %>%
  filter(Measurement=='5')%>%
  dplyr::select(Wavelength, photons)%>%
  rename(Measurement_5 = photons)

# Merge all data frames by Wavelength
no_turb_merged_data <- left_join(no_turb_1, no_turb_2, by = "Wavelength") %>%
  left_join(no_turb_3, by = "Wavelength") %>%
  left_join(no_turb_4, by = "Wavelength") %>%
```

```
  left_join(no_turb_5, by = "Wavelength")
no_turb_merged_data <- no_turb_merged_data %>%
  mutate(
    no_turb_cal_1 = Measurement_5 / Measurement_1,
    no_turb_cal_2 = Measurement_5 / Measurement_2,
    no_turb_cal_3 = Measurement_5 / Measurement_3,
    no_turb_cal_4 = Measurement_5 / Measurement_4,
    no_turb_cal_5 = Measurement_5 / Measurement_5)

# Left join based on matching Wavelength values for each Turbidity_Source
all_spectra_adjusted <- left_join(all_spectra, no_turb_merged_data, by = "Wavelength") %>%
  mutate(
    adjusted_photons = case_when(
      Turbidity_Source == "Bentonite" ~ photons * no_turb_cal_1,
      Turbidity_Source == "Algae" ~ photons * no_turb_cal_5,
      Turbidity_Source == "Calcium.Carbonate" ~ photons * no_turb_cal_2,
      Turbidity_Source == "Kaolin" ~ photons * no_turb_cal_4,
      Turbidity_Source == "Charcoal" ~ photons * no_turb_cal_3,
      TRUE ~ photons))%>%  # Keep original photons if Turbidity_Source doesn't match any of the above
  dplyr::select(-no_turb_cal_1, -no_turb_cal_2, -no_turb_cal_3, -no_turb_cal_4, -no_turb_cal_5)  # Remove no_turb_cal columns
# Now, all_spectra_adjusted contains adjusted_photons column with multiplied values for different Turbidity_Source values
# Reorder levels of Turbidity_Source
all_spectra_adjusted$Turbidity_Source <- factor(all_spectra_adjusted$Turbidity_Source,
                                       levels = c("Algae", "Bentonite", "Calcium.Carbonate", "Kaolin", "Charcoal", "No_Turbidity"))







#########################################################################
#######################################################################
```

```
#------------------------------------------------------------------------------------------------------------------------------------------------#
#                                   Calculate luminance, hue, and chroma
#------------------------------------------------------------------------------------------------------------------------------------------------#
###########################################################################
#########################################################################
summary_spectra <- all_spectra_adjusted %>%
  group_by(Turbidity_Source, Measurement, Average_NTU) %>%
  summarise(
    total_photons = sum(adjusted_photons),
    B = sum(adjusted_photons[Wavelength >= 400 & Wavelength < 475]) / sum(adjusted_photons),
    G = sum(adjusted_photons[Wavelength >= 475 & Wavelength < 550]) / sum(adjusted_photons),
    Y = sum(adjusted_photons[Wavelength >= 550 & Wavelength < 625]) / sum(adjusted_photons),
    R = sum(adjusted_photons[Wavelength >= 625 & Wavelength <= 700]) / sum(adjusted_photons),
    LM = R - G,
    MS = Y - B,
    chroma = sqrt(LM^2 + MS^2),
    hue = asin(MS / chroma))
NT_values <- summary_spectra %>%
  filter(Turbidity_Source == 'No_Turbidity', Measurement == '5') %>%
  summarise(NT_LM = LM, NT_MS = MS, NT_total_photons = total_photons, NT_Average_NTU = Average_NTU, NT_hue = hue, NT_chroma = chroma)



#--------------------------------------------------------------------------------------------------------------------#
#------------------------------------------------- Modelling ------------------------------------------------#
#--------------------------------------------------------------------------------------------------------------------#
summary_spectra2<- summary_spectra[summary_spectra$Turbidity_Source != 'Charcoal',]
#remove all No_Turbidity values where Measurement !=5
summary_spectra2 <- summary_spectra2 %>%
  filter(Turbidity_Source != "No_Turbidity" | Measurement == 5)
```

```
summary_spectra2$Turbidity_Source <- factor(summary_spectra2$Turbidity_Source,
                                  levels = c("No_Turbidity", "Algae", "Bentonite", "Calcium.Carbonate", "Kaolin"))

Exp_design$Michelson_Contrast <- abs(Exp_design$Michelson_Contrast) # absolute values
Exp_design$Turbidity_Source <- factor(Exp_design$Turbidity_Source,
                              levels = c("Baseline", "Algae", "Bentonite", "Calcium Carbonate", "Kaolin"))
Exp_design_1<-Exp_design
Exp_design_2 <- Exp_design_1[!is.na(Exp_design_1$Average_NTU), ] # remove NAs


filtered_data <- Exp_design %>%
  filter(Time %in% c(0, 6))
grouped_data <- filtered_data %>%
  group_by(Turbidity_Source, Date) %>%
  summarize(# Calculate percentage change between Time = 0 and Time = 1
    Change = ((Average_NTU[2] - Average_NTU[1]) / Average_NTU[1]) * 100) # Calculate the mean and standard error of the change
result_summary <- grouped_data %>%
  group_by(Turbidity_Source) %>%
  summarize(
    AvgChange = mean(Change, na.rm = TRUE),
    SEChange = sd(Change, na.rm = TRUE) / sqrt(n()),)  # Standard error
print(result_summary)

# Calculate Percentage_Difference as the percentage difference from Time=0 values
grouped_data <- Exp_design %>%
  filter(Turbidity_Source != "Baseline") %>%
  group_by(Turbidity_Source, Date) %>%
  mutate(Percentage_Difference_SR = (Average_NTU - first(Average_NTU)) / first(Average_NTU) * 100)%>%
  mutate(Percentage_Difference_pH = (pH - first(pH)) / first(pH) * 100)%>%
  mutate(Percentage_Difference_KH = (KH - first(KH)) / first(KH) * 100)

mean_first_values <- Exp_design %>%
  group_by(Turbidity_Source) %>%
```

```
  summarize(mean_first_pH = mean(first(pH)), mean_first_KH = mean(first(KH)))

# Calculate summary statistics for Percentage_Difference
summary_data <- grouped_data %>%
  group_by(Turbidity_Source, Time) %>%
  summarize(
    Mean_Percentage_Difference_SR = mean(Percentage_Difference_SR, na.rm = TRUE),
    SE_Percentage_Difference_SR = sd(Percentage_Difference_SR, na.rm = TRUE) / sqrt(n()),
    Mean_Percentage_Difference_pH = mean(Percentage_Difference_pH, na.rm = TRUE),
    SE_Percentage_Difference_pH = sd(Percentage_Difference_pH, na.rm = TRUE) / sqrt(n()),
    Mean_Percentage_Difference_KH = mean(Percentage_Difference_KH, na.rm = TRUE),
    SE_Percentage_Difference_KH = sd(Percentage_Difference_KH, na.rm = TRUE) / sqrt(n())
  )%>%
  mutate(Percentage_of_Start_SR = Mean_Percentage_Difference_SR + 100)

summary_data <- summary_data %>%
  left_join(mean_first_values, by = "Turbidity_Source") %>%
  mutate(Change_from_Baseline_pH = (Mean_Percentage_Difference_pH / 100 * mean_first_pH) + mean_first_pH,
         Change_from_Baseline_KH = (Mean_Percentage_Difference_KH / 100 * mean_first_KH) + mean_first_KH)




#Luminance
TP9<-lmer(total_photons ~ Average_NTU : Turbidity_Source + I(Average_NTU^ 2) + (1|Turbidity_Source), data=summary_spectra2)#;
simulateResiduals(fittedModel=TP9,plot=T)
summary(TP9)

#Hue:
M3<- lmer(hue ~ Average_NTU : Turbidity_Source + (1|Turbidity_Source), data=summary_spectra2)#; simulateResiduals(fittedModel = M3, plot = TRUE)
```

```
summary(M3)

#Chroma
M3<- lmer(chroma ~ Average_NTU : Turbidity_Source + (1|Turbidity_Source), data=summary_spectra2)#; simulateResiduals(fittedModel = M3, plot = TRUE)
summary(M3)

#Image contrast
M8<- lmer(Michelson_Contrast ~ Average_NTU + Turbidity_Source + I(Average_NTU^ 2) + (1|Turbidity_Source) + (1|Camera_Location), data = Exp_design_1)#; simulateResiduals(fittedModel = M8, plot = TRUE)
summary(M8)

#Setting Rate
Exp_design_2$transformed_log <- log(Exp_design_2$Average_NTU); #hist(Exp_design_2$transformed_log)
M2<- lmer(transformed_log ~ Time : Turbidity_Source + Turbidity_Source + (1|Turbidity_Source) + (1|Starting_Turbidity_level), data = Exp_design_2)#; simulateResiduals(fittedModel = M2, plot = TRUE)
summary(M2)

#pH
M8<- lmer(pH ~ Turbidity_Source : Time + Turbidity_Source + (1|Turbidity_Source)+ (1|Starting_Turbidity_level), data = Exp_design_1)#; simulateResiduals(fittedModel = M8, plot = TRUE)
summary(M8)

#KH
M10<- lmer(KH ~ Turbidity_Source + Time + (1|Turbidity_Source)+ (1|Starting_Turbidity_level), data = Exp_design_1)#; simulateResiduals(fittedModel = M10, plot = TRUE)
summary(M10)



###############################################################################
############################################################################
```

```
#-----------------------------------------------------------------------------------------------------------------
--------------------#
#                                          Separating data for absolute irradiance plots
#-----------------------------------------------------------------------------------------------------------------
--------------------#
##########################################################################
########################################################################

#________________________________          Scaled          Photons          Calculations
________________________________#
# Filter all_spectra_adjusted for Measurement 5 and Turbidity_Source = No_Turbidity
no_turb_5_adjusted <- all_spectra_adjusted %>%
  filter(Measurement == 5, Turbidity_Source == "No_Turbidity") %>%
  dplyr::select(Wavelength, adjusted_photons, Turbidity_Source)
NTU_10 <- all_spectra_adjusted %>% filter(Average_NTU >= 7 & Average_NTU <= 13)%>%
#10 +/- 3
  filter(Turbidity_Source != "Charcoal")
NTU_10 <- bind_rows(NTU_10, no_turb_5_adjusted)
NTU_33 <- all_spectra_adjusted %>% filter(Average_NTU >= 31 & Average_NTU <= 35)
%>% #33 +/- 2
  filter(Turbidity_Source != "Charcoal")
NTU_33 <- bind_rows(NTU_33, no_turb_5_adjusted)
NTU_52 <- all_spectra_adjusted %>% filter(Average_NTU >= 47.75 & Average_NTU <=
56.75)%>% #52.25 +/- 4.5
  filter(Turbidity_Source != "Charcoal")
NTU_52 <- bind_rows(NTU_52, no_turb_5_adjusted)
NTU_94 <- all_spectra_adjusted %>% filter(Average_NTU >= 92 & Average_NTU <= 96)%>%
#94 +/- 2
  filter(Turbidity_Source != "Charcoal")
NTU_94 <- bind_rows(NTU_94, no_turb_5_adjusted)

#10 NTU
# Function to calculate the area under the curve for each Turbidity_Source
calculate_area <- function(data) {
  area <- trapz(data$Wavelength, data$adjusted_photons)
  return(data.frame(Area_Under_Curve = area))}
area_under_curve_data <- NTU_10 %>%
```

```
  group_by(Turbidity_Source) %>%
  nest() %>%
  mutate(area_data = map(data, calculate_area)) %>%
  unnest(cols = c(area_data))
print(area_under_curve_data)

NTU_10 <- NTU_10 %>% #makes the area under each curve equaled to 1
  mutate(standardized_photons = case_when(
    Turbidity_Source == 'Bentonite' ~ adjusted_photons /  2.43e20,
    Turbidity_Source == 'Calcium.Carbonate' ~ adjusted_photons / 2.18e20,
    Turbidity_Source == 'Kaolin' ~ adjusted_photons / 2.46e20,
    Turbidity_Source == 'Algae' ~ adjusted_photons / 9.17e19,
    Turbidity_Source == 'No_Turbidity' ~ adjusted_photons / 1.75e20,
    TRUE ~ adjusted_photons  # For other cases, keep the original value
  ))%>%
  group_by(Wavelength)

calculatescaled_area <- function(data) {
  area <- trapz(data$Wavelength, data$standardized_photons)
  return(data.frame(Area_Under_Curve = area))}

area_under_curve_data <- NTU_10 %>%
  group_by(Turbidity_Source) %>%
  nest() %>%
  mutate(area_data = map(data, calculatescaled_area)) %>%
  unnest(cols = c(area_data))
print(area_under_curve_data)


#33 NTU
area_under_curve_data <- NTU_33 %>%
  group_by(Turbidity_Source) %>%
  nest() %>%
  mutate(area_data = map(data, calculate_area)) %>%
  unnest(cols = c(area_data))
print(area_under_curve_data)
```

```
NTU_33 <- NTU_33 %>%
  mutate(standardized_photons = case_when(
    Turbidity_Source == 'Bentonite' ~ adjusted_photons /  2.47e20,
    Turbidity_Source == 'Calcium.Carbonate' ~ adjusted_photons / 2.64e20,
    Turbidity_Source == 'Kaolin' ~ adjusted_photons / 3.26e20,
    Turbidity_Source == 'Algae' ~ adjusted_photons / 6.86e19,
    Turbidity_Source == 'No_Turbidity' ~ adjusted_photons / 1.75e20,
    TRUE ~ adjusted_photons  # For other cases, keep the original value
  ))%>%
  group_by(Wavelength)

area_under_curve_data <- NTU_33 %>%
  group_by(Turbidity_Source) %>%
  nest() %>%
  mutate(area_data = map(data, calculatescaled_area)) %>%
  unnest(cols = c(area_data))
print(area_under_curve_data)


#52 NTU
area_under_curve_data <- NTU_52 %>%
  group_by(Turbidity_Source) %>%
  nest() %>%
  mutate(area_data = map(data, calculate_area)) %>%
  unnest(cols = c(area_data))
print(area_under_curve_data)

NTU_52 <- NTU_52 %>%
  mutate(standardized_photons = case_when(
    Turbidity_Source == 'Bentonite' ~ adjusted_photons /  2.28e20,
    Turbidity_Source == 'Calcium.Carbonate' ~ adjusted_photons / 2.73e20,
    Turbidity_Source == 'Kaolin' ~ adjusted_photons / 3.42e20,
    Turbidity_Source == 'Algae' ~ adjusted_photons / 5.26e19,
    Turbidity_Source == 'No_Turbidity' ~ adjusted_photons / 1.75e20,
    TRUE ~ adjusted_photons  # For other cases, keep the original value
  ))%>%
  group_by(Wavelength)
```

```
area_under_curve_data <- NTU_52 %>%
  group_by(Turbidity_Source) %>%
  nest() %>%
  mutate(area_data = map(data, calculatescaled_area)) %>%
  unnest(cols = c(area_data))
print(area_under_curve_data)


#94 NTU
area_under_curve_data <- NTU_94 %>%
  group_by(Turbidity_Source) %>%
  nest() %>%
  mutate(area_data = map(data, calculate_area)) %>%
  unnest(cols = c(area_data))
print(area_under_curve_data)

NTU_94 <- NTU_94 %>%
  mutate(standardized_photons = case_when(
    Turbidity_Source == 'Bentonite' ~ adjusted_photons /  1.98e20,
    Turbidity_Source == 'Calcium.Carbonate' ~ adjusted_photons / 2.98e20,
    Turbidity_Source == 'Kaolin' ~ adjusted_photons / 3.53e20,
    Turbidity_Source == 'No_Turbidity' ~ adjusted_photons / 1.75e20,
    TRUE ~ adjusted_photons  # For other cases, keep the original value
  ))%>%
  group_by(Wavelength)

area_under_curve_data <- NTU_94 %>%
  group_by(Turbidity_Source) %>%
  nest() %>%
  mutate(area_data = map(data, calculatescaled_area)) %>%
  unnest(cols = c(area_data))
print(area_under_curve_data)
```

```
##########################################################################
########################################################################
#--------------------------------------------------------------------------------------------------------------
--------------------#
#----------------------------------------------------------------- Plots ----------------------------------------
--------------------#
#--------------------------------------------------------------------------------------------------------------
--------------------#
##########################################################################
########################################################################

#------------------------------------------------------------------------#
#---------------------------- Absolute Irradiance ---------------------------#
#------------------------------------------------------------------------#
# 10 NTU
plot1a <- ggplot(NTU_10, aes(x = Wavelength, y = adjusted_photons, color =
Turbidity_Source)) +
  geom_line(linetype = ifelse(NTU_10$Turbidity_Source == "No_Turbidity", "dashed", "solid"))
+  # Add linetype argument
  labs(x = "Wavelength (nm)", y = bquote("photons cm"^"-2"* s^-1*nm^-1), color = "Turbidity
Source", title = "10 NTU") +
  theme_classic() +
  scale_color_manual(name = "Turbidity Source",
                     values = c("Algae" = "#90d743", "Bentonite" = "#21918c", "Calcium.Carbonate"
= "#3b528b", "Kaolin" = "#440154", "No_Turbidity" = "red"),
                     labels = c("Algae", "Bentonite", "Calcium Carbonate", "Kaolin", "No Turbidity"),
                     guide = guide_legend()) +
  scale_y_continuous(limits = c(0, 3.5e+18))
plot1a
plot1b <- ggplot(NTU_10, aes(x = Wavelength, y = standardized_photons, color =
Turbidity_Source)) +
  geom_line(linetype = ifelse(NTU_10$Turbidity_Source == "No_Turbidity", "dashed", "solid"))
+  # Add linetype argument
  labs(x = "Wavelength (nm)", y = bquote("Standardised photons cm"^"-2"* s^-1*nm^-1), color
= "Turbidity Source", title = "10 NTU") +
  theme_classic() +
  scale_color_manual(name = "Turbidity Source",
```

```
                    values = c("Algae" = "#90d743", "Bentonite" = "#21918c", "Calcium.Carbonate"
= "#3b528b", "Kaolin" = "#440154", "No_Turbidity" = "red"),
                    labels = c("Algae", "Bentonite", "Calcium Carbonate", "Kaolin", "No Turbidity"),
                    guide = guide_legend()) +
  scale_y_continuous(limits = c(0, 0.01), labels = scientific_format())

# 33 NTU
plot2a <- ggplot(NTU_33, aes(x = Wavelength, y = adjusted_photons, color =
Turbidity_Source)) +
  geom_line(linetype = ifelse(NTU_33$Turbidity_Source == "No_Turbidity", "dashed", "solid"))
+  # Add linetype argument
  labs(x = "Wavelength (nm)", y = bquote("photons cm"^"-2"* s^-1*nm^-1), color = "Turbidity
Source", title = "33 NTU") +
  theme_classic() +
  scale_color_manual(name = "Turbidity Source",
                    values = c("Algae" = "#90d743", "Bentonite" = "#21918c", "Calcium.Carbonate"
= "#3b528b", "Kaolin" = "#440154", "No_Turbidity" = "red"),
                    labels = c("Algae", "Bentonite", "Calcium Carbonate", "Kaolin", "No Turbidity"),
                    guide = guide_legend()) +
  scale_y_continuous(limits = c(0, 3.5e+18))

plot2b <- ggplot(NTU_33, aes(x = Wavelength, y = standardized_photons, color =
Turbidity_Source)) +
  geom_line(linetype = ifelse(NTU_33$Turbidity_Source == "No_Turbidity", "dashed", "solid"))
+  # Add linetype argument
  labs(x = "Wavelength (nm)", y =  bquote("Standardised photons cm"^"-2"* s^-1*nm^-1), color
= "Turbidity Source", title = "33 NTU") +
  theme_classic() +
  scale_color_manual(name = "Turbidity Source",
                    values = c("Algae" = "#90d743", "Bentonite" = "#21918c", "Calcium.Carbonate"
= "#3b528b", "Kaolin" = "#440154", "No_Turbidity" = "red"),
                    labels = c("Algae", "Bentonite", "Calcium Carbonate", "Kaolin", "No Turbidity"),
                    guide = guide_legend()) +
  scale_y_continuous(limits = c(0, 0.01), labels = scientific_format())

# 52 NTU
```

```
plot3a <- ggplot(NTU_52, aes(x = Wavelength, y = adjusted_photons, color =
Turbidity_Source)) +
  geom_line(linetype = ifelse(NTU_52$Turbidity_Source == "No_Turbidity", "dashed", "solid"))
+  # Add linetype argument
  labs(x = "Wavelength (nm)", y = bquote("photons cm"^"-2"* s^-1*nm^-1), color = "Turbidity
Source", title = "52 NTU") +
  theme_classic() +
  scale_color_manual(name = "Turbidity Source",
                     values = c("Algae" = "#90d743", "Bentonite" = "#21918c", "Calcium.Carbonate"
= "#3b528b", "Kaolin" = "#440154", "No_Turbidity" = "red"),
                     labels = c("Algae", "Bentonite", "Calcium Carbonate", "Kaolin", "No Turbidity"),
                     guide = guide_legend()) +
  scale_y_continuous(limits = c(0, 3.5e+18))

plot3b <- ggplot(NTU_52, aes(x = Wavelength, y = standardized_photons, color =
Turbidity_Source)) +
  geom_line(linetype = ifelse(NTU_52$Turbidity_Source == "No_Turbidity", "dashed", "solid"))
+  # Add linetype argument
  labs(x = "Wavelength (nm)", y =  bquote("Standardised photons cm"^"-2"* s^-1*nm^-1), color
= "Turbidity Source", title = "52 NTU") +
  theme_classic() +
  scale_color_manual(name = "Turbidity Source",
                     values = c("Algae" = "#90d743", "Bentonite" = "#21918c", "Calcium.Carbonate"
= "#3b528b", "Kaolin" = "#440154", "No_Turbidity" = "red"),
                     labels = c("Algae", "Bentonite", "Calcium Carbonate", "Kaolin", "No Turbidity"),
                     guide = guide_legend()) +
  scale_y_continuous(limits = c(0, 0.01), labels = scientific_format())

# 94 NTU
plot4a <- ggplot(NTU_94, aes(x = Wavelength, y = adjusted_photons, color =
Turbidity_Source)) +
  geom_line(linetype = ifelse(NTU_94$Turbidity_Source == "No_Turbidity", "dashed", "solid"))
+  # Add linetype argument
  labs(x = "Wavelength (nm)", y = bquote("photons cm"^"-2"* s^-1*nm^-1), color = "Turbidity
Source", title = "94 NTU") +
  theme_classic() +
  scale_color_manual(name = "Turbidity Source",
```

```
                    values = c("Algae" = "#90d743", "Bentonite" = "#21918c", "Calcium.Carbonate"
= "#3b528b", "Kaolin" = "#440154", "No_Turbidity" = "red"),
                    labels = c("Bentonite", "Calcium Carbonate", "Kaolin", "No Turbidity"),
                    guide = guide_legend()) +
  scale_y_continuous(limits = c(0, 3.5e+18))

plot4b <- ggplot(NTU_94, aes(x = Wavelength, y = standardized_photons, color =
Turbidity_Source)) +
  geom_line(linetype = ifelse(NTU_94$Turbidity_Source == "No_Turbidity", "dashed", "solid"))
+  # Add linetype argument
  labs(x = "Wavelength (nm)", y =  bquote("Standardised photons cm"^"-2"* s^-1*nm^-1), color
= "Turbidity Source", title = "94 NTU") +
  theme_classic() +
  scale_color_manual(name = "Turbidity Source",
                    values = c("Algae" = "#90d743", "Bentonite" = "#21918c", "Calcium.Carbonate"
= "#3b528b", "Kaolin" = "#440154", "No_Turbidity" = "red"),
                    labels = c("Bentonite", "Calcium Carbonate", "Kaolin", "No Turbidity"),
                    guide = guide_legend()) +
  scale_y_continuous(limits = c(0, 0.01), labels = scientific_format())


#--------------------------------------------------------------------------#
#---------------      Luminance, hue and chroma plots        -----------------#
#--------------------------------------------------------------------------#
summary_spectra1 <- summary_spectra[summary_spectra$Turbidity_Source !=
'No_Turbidity',]
summary_spectra1$Turbidity_Source <- factor(summary_spectra1$Turbidity_Source,
                                    levels = c("Algae", "Bentonite", "Calcium.Carbonate", "Kaolin",
"Charcoal"))
summary_spectra2<- summary_spectra1[summary_spectra1$Turbidity_Source !=
'Charcoal',]
plot1<-ggplot(summary_spectra2, aes(x = Average_NTU, y = total_photons, color =
Turbidity_Source, shape=Turbidity_Source)) +
  geom_path(aes(group = Turbidity_Source), na.rm = TRUE) +
  geom_point() +
  #geom_point(data = NT_values, aes(x = NT_Average_NTU, y = NT_total_photons), color =
"red")+
```

```
  geom_hline(yintercept = NT_values$NT_total_photons, linetype = "dashed", color = "red") +
  labs(x = "Turbidity Level (NTU)", y = bquote("Luminance (photons cm"^"-2"* "s"^"-1"*"nm"^"-
1")) +
  theme_classic() +
  scale_color_manual(name = "Turbidity Source",
                     values = c("Algae" = "#90d743", "Bentonite" = "#21918c", "Calcium.Carbonate"
= "#3b528b", "Kaolin" = "#440154","Charcoal" = "#fde725", "No Turbidity" = "red"),
                     labels = c("Algae", "Bentonite", "Calcium Carbonate", "Kaolin", "Charcoal"),
                     guide = guide_legend())+
  scale_shape_manual(name = "Turbidity Source",
                     values = c("Algae" = 18, "Bentonite" = 15, "Calcium.Carbonate" = 16, "Kaolin" =
17, "Charcoal"=18),
                     labels = c("Algae", "Bentonite", "Calcium Carbonate", "Kaolin", "Charcoal"),
                     guide = guide_legend())
plot1

plot2<-ggplot(summary_spectra2, aes(x = Average_NTU, y = hue, color = Turbidity_Source,
shape=Turbidity_Source)) +
  geom_point() +
  geom_path(aes(group = Turbidity_Source), na.rm = TRUE) +
  #geom_point(data = NT_values, aes(x = NT_Average_NTU, y = NT_hue), color = "red")+
  geom_hline(yintercept = NT_values$NT_hue, linetype = "dashed", color = "red") +
  labs(x = "Turbidity Level (NTU)", y = "Hue") +
  theme_classic()+
  scale_color_manual(name = "Turbidity Source",
                     values = c("Algae" = "#90d743", "Bentonite" = "#21918c", "Calcium.Carbonate"
= "#3b528b", "Kaolin" = "#440154","Charcoal" = "#fde725", "No Turbidity" = "red"),
                     labels = c("Algae", "Bentonite", "Calcium Carbonate", "Kaolin", "Charcoal"),
                     guide = guide_legend())+
  scale_shape_manual(name = "Turbidity Source",
                     values = c("Algae" = 18, "Bentonite" = 15, "Calcium.Carbonate" = 16, "Kaolin" =
17),
                     labels = c("Algae", "Bentonite", "Calcium Carbonate", "Kaolin"),
                     guide = guide_legend())

plot3<-ggplot(summary_spectra2, aes(x = Average_NTU, y = chroma, color =
Turbidity_Source, shape=Turbidity_Source)) +
```

```
  geom_path(aes(group = Turbidity_Source), na.rm = TRUE) +
  geom_point() +
  geom_point(data = NT_values, aes(x = NT_Average_NTU, y = NT_chroma), color = "red")+
  geom_hline(yintercept = NT_values$NT_chroma, linetype = "dashed", color = "red") +
  labs(x = "Turbidity Level (NTU)", y = "Chroma") +
  theme_classic()+
  scale_color_manual(name = "Turbidity Source",
                     values = c("Algae" = "#90d743", "Bentonite" = "#21918c", "Calcium.Carbonate"
= "#3b528b", "Kaolin" = "#440154","Charcoal" = "#fde725", "No Turbidity" = "red"),
                     labels = c("Algae", "Bentonite", "Calcium Carbonate", "Kaolin", "Charcoal"),
                     guide = guide_legend())+
  scale_shape_manual(name = "Turbidity Source",
                     values = c("Algae" = 18, "Bentonite" = 15, "Calcium.Carbonate" = 16, "Kaolin" =
17),
                     labels = c("Algae", "Bentonite", "Calcium Carbonate", "Kaolin"),
                     guide = guide_legend())


#------------------------------------#
#----------- Image Contrast -----------#
#-------------------------------------#
IC_plot_data <- Exp_design_1 %>%
  filter(Average_NTU < 51)
plot4<-ggplot(IC_plot_data, aes(x = Average_NTU, y = Michelson_Contrast, group=
Turbidity_Source, shape=Turbidity_Source)) +
  geom_point(aes(color = Turbidity_Source)) +
  geom_smooth(method = "lm", formula = y ~ x + I(x^2), aes(color=Turbidity_Source), se =
FALSE, size = 0.5) +  # Add the regression line
  geom_hline(yintercept = baseline_avgIC, linetype = "dashed", color = "red") +
  labs(x = "Turbidity Level (NTU)", y = "Image Contrast") +
  theme_classic() +
  scale_color_manual(name = "Turbidity Source",
                     values = c("Algae" = "#90d743", "Bentonite" = "#21918c", "Calcium Carbonate"
= "#3b528b", "Kaolin" = "#440154"),
                     labels = c("Algae", "Bentonite", "Calcium Carbonate", "Kaolin"),
                     guide = guide_legend())+
  scale_shape_manual(name = "Turbidity Source",
```

```
                    values = c("Algae" = 18, "Bentonite" = 15, "Calcium Carbonate" = 16, "Kaolin" =
17),
                    labels = c("Algae", "Bentonite", "Calcium Carbonate", "Kaolin"),
                    guide = guide_legend())
plot4

#------------------------------------#
#---------- Settling Rate -------------#
#------------------------------------#
plot5<-ggplot(summary_data, aes(x = Time, y = Percentage_of_Start_SR, group =
Turbidity_Source, shape = Turbidity_Source, color = Turbidity_Source)) +
  geom_point() +
  geom_line()+
  geom_hline(yintercept = baseline_avgSR, linetype = "dashed", color = "red") +
  labs(x = "Time (hours)", y = "Turbidity Level (NTU)") +
  theme_classic() +
  scale_color_manual(name = "Turbidity Source",
                    values = c("Algae" = "#90d743", "Bentonite" = "#21918c", "Calcium Carbonate"
= "#3b528b", "Kaolin" = "#440154"),
                    labels = c("Algae", "Bentonite", "Calcium Carbonate", "Kaolin"),
                    guide = guide_legend()) +
  scale_shape_manual(name = "Turbidity Source",
                    values = c("Algae" = 18, "Bentonite" = 15, "Calcium Carbonate" = 16, "Kaolin" =
17),
                    labels = c("Algae", "Bentonite", "Calcium Carbonate", "Kaolin"),
                    guide = guide_legend())+
  scale_y_continuous(limits = c(0, 100))  # Set y-axis range from 0 to 100
plot5

#------------------------------------#
#---------------- pH ----------------#
#------------------------------------#
summary_data_no_na                                                                    <-
summary_data[complete.cases(summary_data$Change_from_Baseline_KH), ]
plot6<-ggplot(summary_data_no_na, aes(x = Time, y = Change_from_Baseline_pH, group =
Turbidity_Source, shape = Turbidity_Source, color = Turbidity_Source)) +
  geom_point() +
```

```
  geom_hline(yintercept = baseline_avgpH, linetype = "dashed", color = "red") +
  geom_line()+
  labs(x = "Time (hours)", y = "pH") +
  theme_classic() +
  scale_color_manual(name = "Turbidity Source",
                     values = c("Algae" = "#90d743", "Bentonite" = "#21918c", "Calcium Carbonate"
= "#3b528b", "Kaolin" = "#440154"),
                     labels = c("Algae", "Bentonite", "Calcium Carbonate", "Kaolin"),
                     guide = guide_legend()) +
  scale_shape_manual(name = "Turbidity Source",
                     values = c("Algae" = 18, "Bentonite" = 15, "Calcium Carbonate" = 16, "Kaolin" =
17),
                     labels = c("Algae", "Bentonite", "Calcium Carbonate", "Kaolin"),
                     guide = guide_legend())
plot6

#------------------------------------#
#---------------- KH ----------------#
#------------------------------------#
plot7<-ggplot(summary_data_no_na, aes(x = Time, y = Change_from_Baseline_KH, group =
Turbidity_Source, shape = Turbidity_Source, color = Turbidity_Source)) +
  geom_point() +
  geom_line()+
  geom_hline(yintercept = baseline_avgKH, linetype = "dashed", color = "red") +
  labs(x = "Time (hours)", y = "KH") +
  theme_classic() +
  scale_color_manual(name = "Turbidity Source",
                     values = c("Algae" = "#90d743", "Bentonite" = "#21918c", "Calcium Carbonate"
= "#3b528b", "Kaolin" = "#440154"),
                     labels = c("Algae", "Bentonite", "Calcium Carbonate", "Kaolin"),
                     guide = guide_legend()) +
  scale_shape_manual(name = "Turbidity Source",
                     values = c("Algae" = 18, "Bentonite" = 15, "Calcium Carbonate" = 16, "Kaolin" =
17),
                     labels = c("Algae", "Bentonite", "Calcium Carbonate", "Kaolin"),
                     guide = guide_legend())+
  theme(plot.title = element_text(size = 17))  # Adjust the size as needed
```

```
plot7



#--------------------------------------------------------------------------#
#--------------------------    Composite Plot     ---------------------------#
#--------------------------------------------------------------------------#

# Remove legends from each plot
plot1a <- plot1a + theme(legend.position = "none")
plot1b <- plot1b + theme(legend.position = "none")
plot2a <- plot2a + theme(legend.position = "none")
plot2b <- plot2b + theme(legend.position = "none")
plot3a <- plot3a + theme(legend.position = "none")
plot3b <- plot3b + theme(legend.position = "none")
plot4a <- plot4a + theme(legend.position = "none")
plot4b <- plot4b + theme(legend.position = "none")
plot1 <- plot1 + theme(legend.position = "none")
plot2 <- plot2 + theme(legend.position = "none")
plot3 <- plot3 + theme(legend.position = "none")
plot4 <- plot4 + theme(legend.position = "none")
plot5 <- plot5 + theme(legend.position = "none")
plot6 <- plot6 + theme(legend.position = "none")
plot7 <- plot7 + theme(legend.position = "none")


composite_plot <- grid.arrange(
  plot1a + theme(plot.title = element_text(hjust = 0.5, size = 10)),
  plot2a + theme(plot.title = element_text(hjust = 0.5, size = 10)),
  plot3a + theme(plot.title = element_text(hjust = 0.5, size = 10)),
  plot4a + theme(plot.title = element_text(hjust = 0.5, size = 10)),
  plot1b + theme(plot.title = element_text(hjust = 0.5, size = 10)),
  plot2b + theme(plot.title = element_text(hjust = 0.5, size = 10)),
  plot3b + theme(plot.title = element_text(hjust = 0.5, size = 10)),
  plot4b + theme(plot.title = element_text(hjust = 0.5, size = 10)),
  plot1 + theme(plot.title = element_text(hjust = 0.5, size = 10)),
  plot2 + theme(plot.title = element_text(hjust = 0.5, size = 10)),
```

```
plot3 + theme(plot.title = element_text(hjust = 0.5, size = 10)),
plot4 + theme(plot.title = element_text(hjust = 0.5, size = 10)),
plot5 + theme(plot.title = element_text(hjust = 0.5, size = 10)),
plot6 + theme(plot.title = element_text(hjust = 0.5, size = 10)),
plot7 + theme(plot.title = element_text(hjust = 0.5, size = 10)),
ncol = 4)
```
